# Novel age-dependent cortico-subcortical morphologic interactions predict fluid intelligence: A multi-cohort geometric deep learning study

**DOI:** 10.1101/2020.10.14.331199

**Authors:** Yunan Wu, Pierre Besson, Emanuel A. Azcona, S. Kathleen Bandt, Todd B Parrish, Hans C Breiter, Aggelos K. Katsaggelos

## Abstract

Brain structure is tightly coupled with brain functions, but it remains unclear how cognition is related to brain morphology, and what is consistent across neurodevelopment. In this work, we developed graph convolutional neural networks (gCNNs) to predict Fluid Intelligence (Gf) from shapes of cortical ribbons and subcortical structures. T1-weighted MRIs from two independent cohorts, the Human Connectome Project (HCP; age: 28.81±3.70) and the Adolescent Brain Cognitive Development Study (ABCD; age: 9.93±0.62) were independently analyzed. Cortical and subcortical surfaces were extracted and modeled as surface meshes. Three gCNNs were trained and evaluated using six-fold nested cross-validation. Overall, combining cortical and subcortical surfaces yielded the best predictions on both HCP (R=0.454) and ABCD datasets (R=0.314), and outperformed the current literature. Across both datasets, the morphometry of the amygdala and hippocampus, along with temporal, parietal and cingulate cortex consistently drove the prediction of Gf, suggesting a novel reframing of the morphometry underlying Gf.

## Introduction

Pioneering work by Binet and Simon suggested that individuals with higher intelligence learn more quickly (faster reaction time) and effectively (higher accuracy) across a broad array of tasks, from naming objects to defining words, drawing pictures, and solving analogies (*1, 2*). Spearman synthesized these observations into the hypothesis of a generalized intelligence factor (g) which reflects abstract thinking and includes the ability to acquire knowledge, adapt to novelty, develop abstract models, and benefit from schooling and experience (*3, 4*). Further work by Cattell (*5*) split g into fluid intelligence (Gf), which is the capacity to solve novel problems and abstract reasoning, and crystallized intelligence (Gc) which relates to accumulated knowledge (*6–8*). Although Gc and Gf are related and rapidly develop in childhood until adolescence, Gf reaches its steady state prior to a delayed declination whereas Gc continues growing throughout the lifespan (*9, 10*). Of these, Gf has been shown to positively correlate with a vast number of cognitive activities, and to be an important predictor of both educational and professional success (*11*), raising questions as to how to optimally predict it in the context of development and senescence for educational and health optimization purposes, and how to do so given the variability of contributions to Gf from different regions of the brain, including subcortical grey matter regions of the medial temporal lobe and basal ganglia that are largely ignored in the literature but are fundamental for working memory which is associated with Gf, and judgment and decision-making which relates to the use of abstract modeling of value (*12, 13*).

Recent work has sought to understand the neural substrates of Gf as one approach to predicting and calibrating it. This work has focused on a broad array of neuroimaging modalities and lesion models, each of which has its limitations. Studies with functional imaging of cognitive tasks, or of synchrony between resting state oscillations in blood-oxygen level dependent (BOLD) signal, have tended to focus on fronto-parietal networks responsible for integrating sensory and executive functions (*14*) in the form of the parieto-frontal integration theory (P-FIT) (*15*), or combinations of lesion and imaging work (*16*) to explore multiple demand (MD) system contributions to Gf (*17*). In the context of structural brain imaging (i.e., morphometry), studies have evaluated the correlation between brain size and Gf (*18*), or evaluated the contribution of specific cortical areas and white matter fiber bundles to Gf. This research has identified associations between Gf and cortical morphology such as cortical thickness, cortical area, cortical volume, gyrification and grey matter density (*19–21*). Individual cortical morphological metrics have shown rather modest correlations with Gf, and cortical morphometric features correlating with Gf have shown moderate overlap as illustrated by Tadayon and his colleagues. (*21*). Although this study reports a correlation between the shape of basal ganglia structures and Gf (*22*), the relative contribution of subcortical structures involved with emotional memory (e.g., amygdala) and judgment and decision making components of the basal ganglia (e.g., nucleus accumbens) have not been investigated, nor have they been evaluated against cortical contributions, which include areas outside of fronto-parietal networks, such as the temporal cortex (*23*).

Multiple approaches exist for assessing grey matter brain structure: (1) volume (regional volumes of deep grey matter structures) to investigate gross volumetric differences, (2) grey matter density using voxel-based morphometry, (3) shape deformation (surface topology) to investigate localized shape differences. Of these approaches, shape analyses allow for a comparison of surface geometrical properties of structures between groups or with behavior (*24*) that may not have had an overall volume change or alteration in grey matter density, and thus may be very sensitive to subtle changes in their relationship to behavior, diagnosis and development (*25, 26*). For instance, exposure-dependent deformations may precede more pronounced volumetric changes with illicit drug exposure (*27*). Other work has shown that subcortical deformations have been reported in thalami of patients with schizophrenia (*28–31*), obsessive-compulsive disorders (*31*), Parkinson’s disease (*32*), and Tourette’s syndrome (*33*). Of surface geometrical measures, thickness is a topographical measure that is an indicator of the integrity of cytoarchitecture in the cortex (*34*), and of all the topographical measures that can be made, cortical thickness is the most invariant brain-size parameter across mammalian evolution (*35, 36*). Neocortical enlargement depends primarily on growth of surface area (*35, 37, 38*), which thus makes cortical surface measures important in considering similarities across cohorts with significant differences in mean age, if one is going to identify consistent features of brain morphometry related to Gf.

Given these considerations, and the dearth of research on (i) both deep grey matter contributions and cortical contributions to Gf, (ii) absence of work of what is common across disparate age groups, the focus of our work was three-fold. First, we aimed to identify the most predictive brain morphometric features of Gf. Due to the challenges inherent in modeling all the relevant cortical morphologic features and the limited predictive power of individual cortical morphologic features, we used a data-driven approach capable of identifying complex non-linear relationships, potentially across remote brain regions, and implicitly encompassing multiple morphometric features such as cortical thickness, cortical area and gyrification, as well the shape of subcortical structures. The second aim of our study was to assess the contribution of the subcortical structures to Gf either alone or combined with cortical morphology. The third aim specifically focused on investigating how age might be involved in the prediction of Gf. For these purposes, we developed a novel geometric deep learning method capable of extracting relevant cortical and subcortical morphological features (*39*). Our method was data-driven and relied on cortical and subcortical surface models, extracted from automated analysis pipelines, as an input for a graph convolutional neural network (gCNN) to infer Gf. Using a 6-fold cross-validation scheme and two large independent datasets, we evaluated the robustness of our method and the reproducibility of the predictions across two different age groups. Finally, a gradient-based backpropagation method allowed us to map the most predictive cortical and subcortical regions involved in the correct prediction of Gf.

## Results

### HCP Dataset Fluid Intelligence Prediction

Table 1 summarizes the comparative performance of each of the three proposed models used to predict fluid intelligence on the HCP testing dataset across all six folds. Fig. 1 shows the distribution of predictions for each model. All three models were able to successfully predict fluid intelligence scores. However, use of both cortical and subcortical surfaces together achieved the best performance (MSE = 0.834, R = 0.454, *p* = 6.2 × 10^−57^), followed by using only the cortical surface data (MSE = 0.886, R = 0.381, *p* = 2.6 × 10^−39^) with use of subcortical surface data alone producing the least accurate results (MSE = 1.014, R = 0.155, *p* = 2.3 × 10^−8^).

**Fig. 1.**
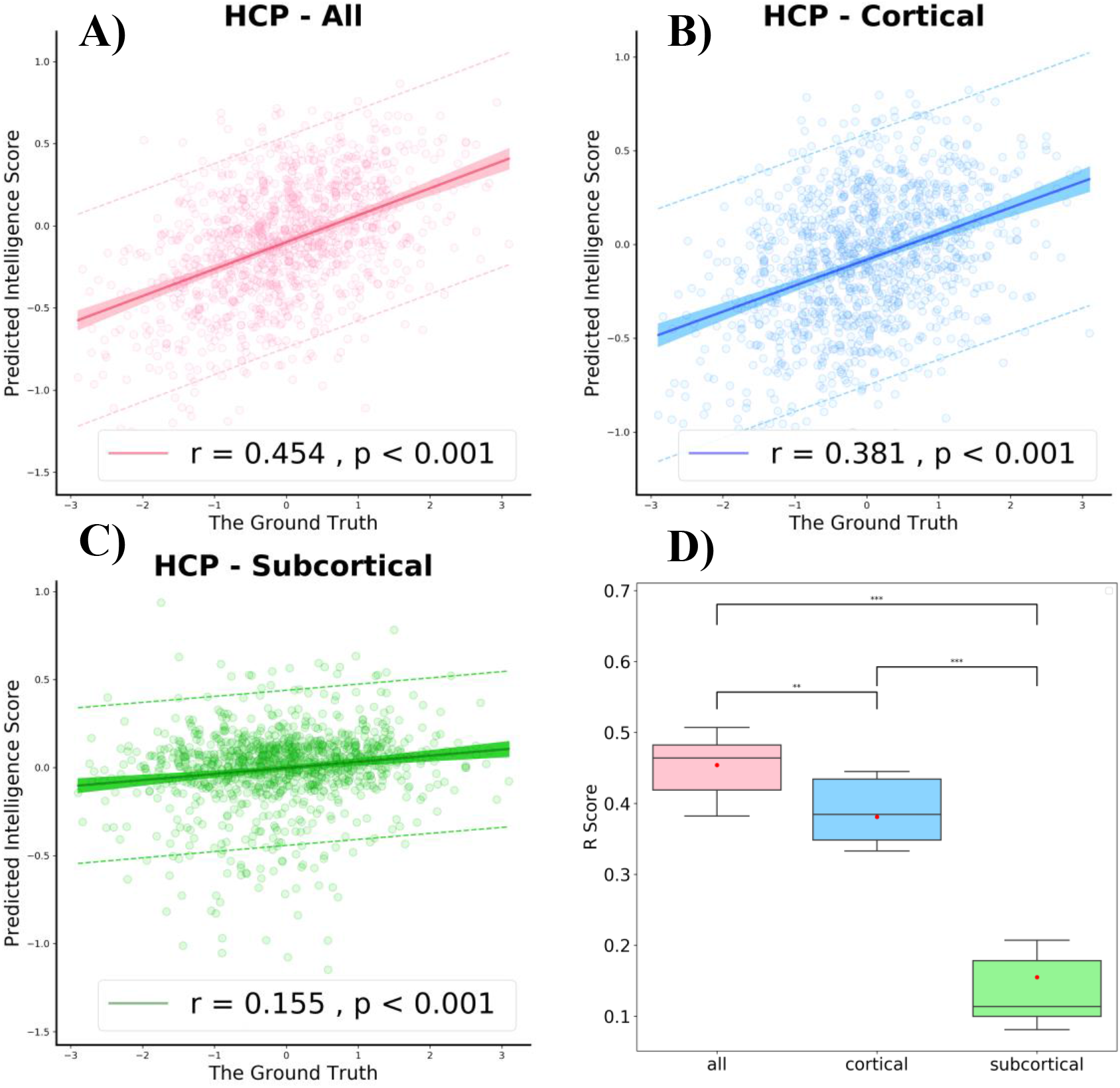
Performance of fluid intelligence score on HCP dataset. A,B,C) prospective predictions on testing dataset using all structures, cortical only and subcortical only respectively. The mean absolute error (MAE) of the prediction is shown by the dashed line. The shaded regions imply the 95% confidence intervals for the regression predictions. The correlation value (R) and p-value of the predicted score vs. the ground truth scores are given. D) comparative boxplots of the R scores over all three different inputs using results across all five folds. The red dots correspond to the R score generated from all testing dataset. (n.s.) Non significant, * p < 0.05, ** p < 0.01, *** p < 0.001.

**Table 1.** Model performance on HCP dataset. The models were trained with sixfold nested cross-validation and the predictions were evaluated on the outer testing set of each fold.

| HCP dataset | Testing Set |  |  |  |  |  |  |  |  |
| --- | --- | --- | --- | --- | --- | --- | --- | --- | --- |
|  | All <sup>a</sup> |  |  | Cor <sup>b</sup> |  |  | Sub <sup>c</sup> |  |  |
|  | R | MSE | Time(s) | R | MSE | Time(s) | R | MSE | Time(s) |
| Fold 1 (N=183) | 0.507 | 0.809 | 1969 | 0.445 | 0.870 | 1084 | 0.197 | 1.071 | 540 |
| Fold 2 (N=183) | 0.465 | 0.915 | 1763 | 0.405 | 0.933 | 997 | 0.122 | 1.055 | 552 |
| Fold 3 (N=183) | 0.463 | 0.889 | 1856 | 0.364 | 0.921 | 1165 | 0.105 | 1.140 | 531 |
| Fold 4 (N=183) | 0.382 | 0.811 | 2041 | 0.343 | 1.033 | 1132 | 0.081 | 0.852 | 490 |
| Fold 5 (N=183) | 0.404 | 0.906 | 1974 | 0.333 | 0.774 | 1057 | 0.098 | 0.938 | 558 |
| Fold 6 (N=182) | 0.488 | 0.677 | 2845 | 0.444 | 0.786 | 1135 | 0.207 | 1.030 | 533 |
| Mean | **0.454** | **0.834** | **1908** | **0.381** | **0.886** | **1095** | **0.155** | **1.014** | **534** |
| ± | ± | ± | ± | ± | ± | ± | ± | ± | ± |
| Sd <sup>d</sup> | **0.049** | **0.090** | **390** | **0.050** | **0.098** | **62** | **0.054** | **0.103** | **24** |

<sup>a</sup>. All: use both cortical and subcortical nodes. <sup>b</sup>. Cor: cortical nodes only. <sup>c</sup>. Sub: subcortical nodes only. <sup>d</sup>. Sd: Standard deviation.

### ABCD Dataset Fluid Intelligence Prediction

Table 2 and Fig. 2 demonstrate the prediction performance for fluid intelligence score using the ABCD testing dataset. Similar to findings from the HCP dataset, performance was significantly improved when combining surface data from both cortical and subcortical surfaces (MSE = 0.919, R = 0.314 *p* = 1.5 × 10^−183^) when using only cortical surface data (MSE = 0.927, R = 0.303, *p* = 7.9 × 10^−171^) or subcortical surface data (MSE = 0.947, R = 0.265, *p* = 4.8 × 10^−130^). Interestingly, the overall performance for fluid intelligence prediction was better on the HCP dataset compared with ABCD dataset.

**Fig. 2.**
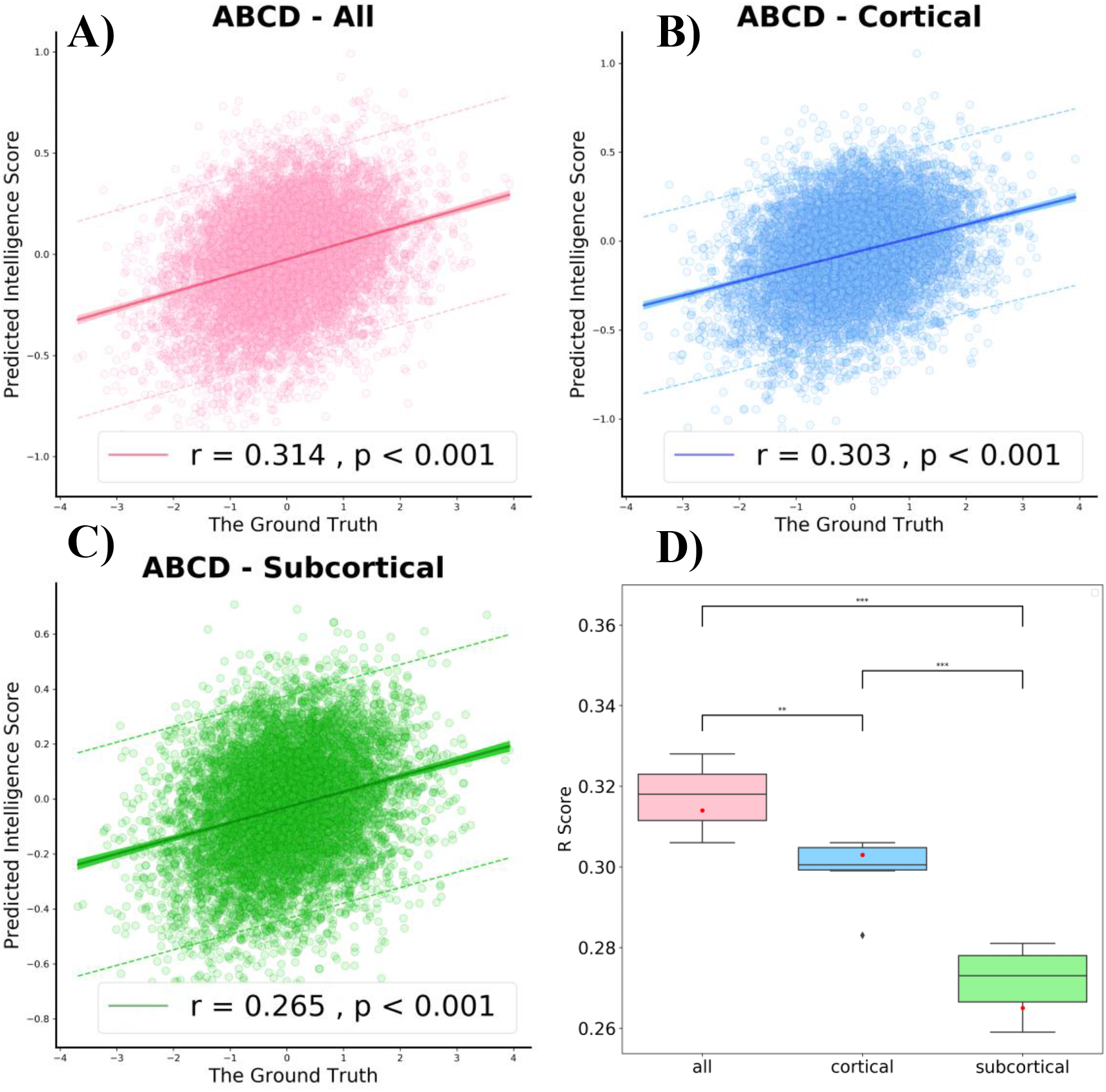
Performance of fluid intelligence score on ABCD dataset. A,B,C) prospective predictions on testing dataset using all structures, cortical only and subcortical only respectively. The mean absolute error (MAE) of the prediction is shown by the dashed line. The shaded regions imply the 95% confidence intervals for the regression predictions. The correlation value (R) and p-value of the predicted score vs. the ground truth scores are given. D) comparative boxplots of the R scores over all three different inputs using results across all five folds. The red dots correspond to the R score generated from all testing dataset. (n.s.) Non significant, * p < 0.05, ** p < 0.01, *** p < 0.001.

**Table 2.**
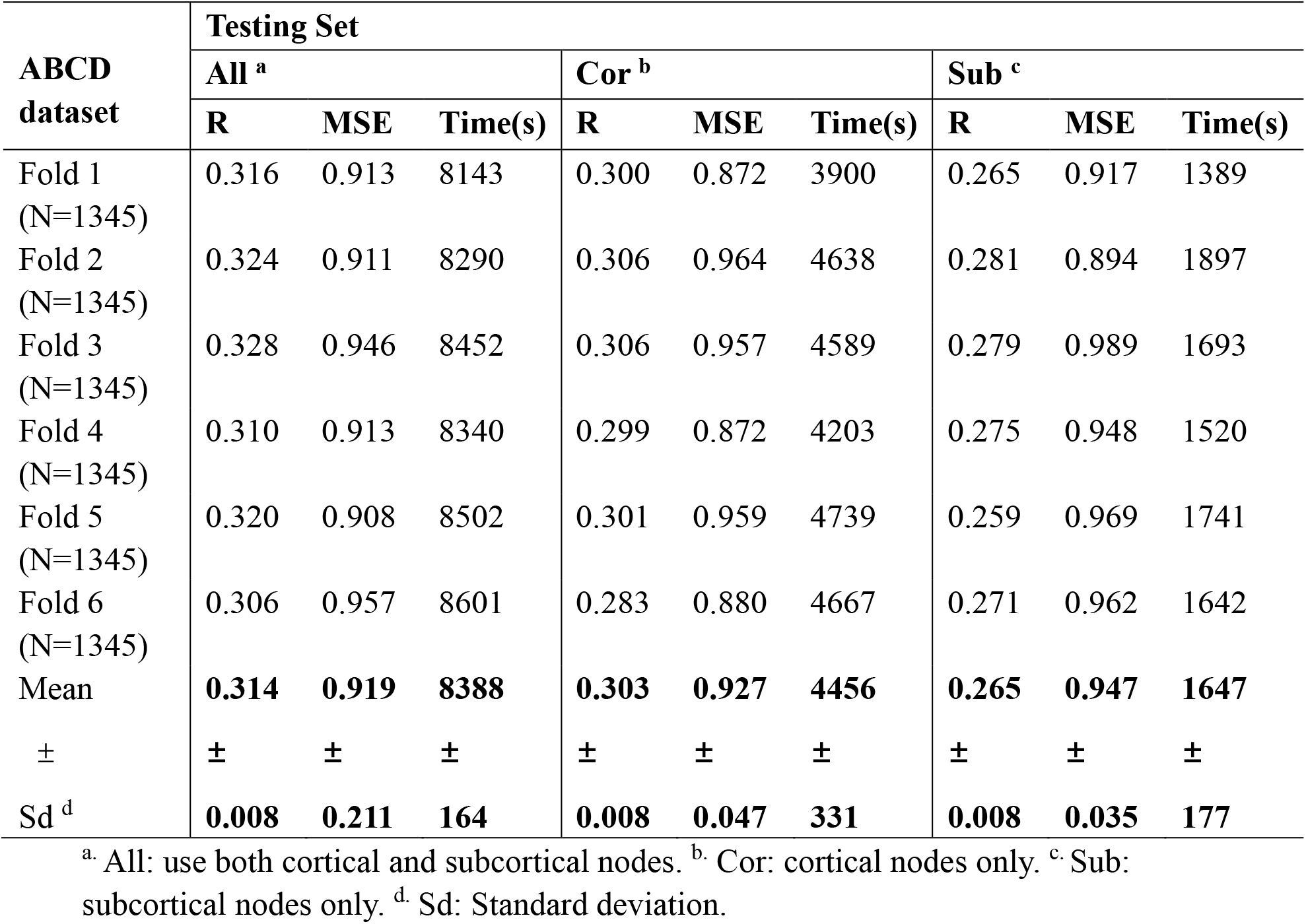
Model performance on ABCD dataset. The models were trained with sixfold nested cross-validation and the predictions were evaluated on the outer testing set of each fold.

Since we used six-fold cross-validation, we had six separate testing datasets and in total 15 correlations were calculated respectively for each inner-fold input and two correlations were calculated inter-cohort. More details can be found in Fig. S1. Table S1 showed the averaged correlations. In each inner-fold, cortical structures showed higher correlations than subcortical structures on both datasets. The correlations on inner-fold inputs in each dataset were higher than those on the inter-cohort correlations across two datasets.

### Mapping Interpretation

In order to provide some interpretability to our models’ performance, we applied a gradient backpropagation-based visualization method (grad-CAM) to visualize the brain areas most relevant to fluid intelligence prediction. Fig. 3 and Fig. 4 show the average maps of the testing sets from both the HCP and ABCD datasets.

**Fig. 3.**
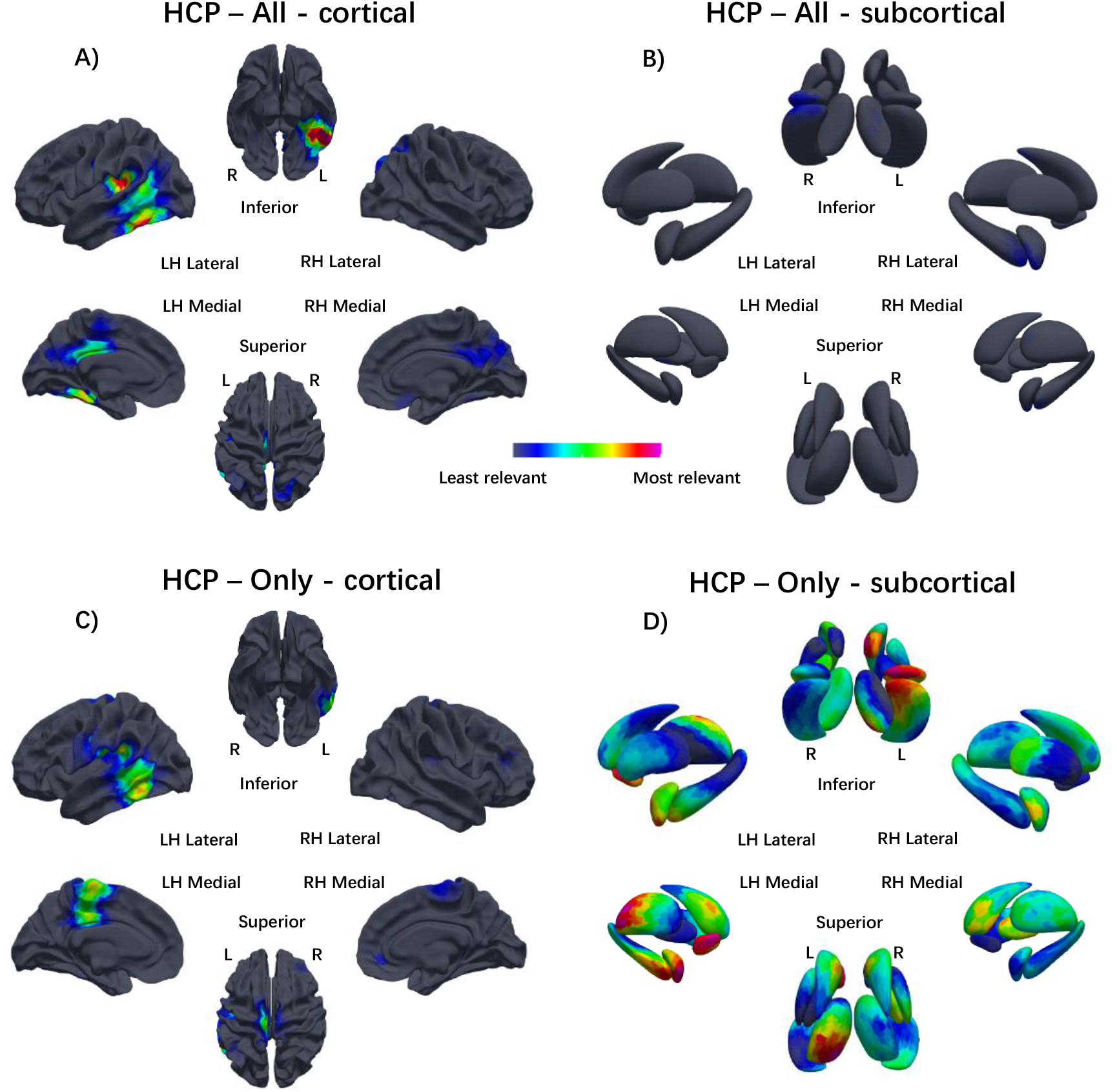
Grad-CAM visualizations on HCP dataset using both cortical and subcortical structures as input. Brain regions predictve of fluid intelligence. Network trained on all cortical and subcortical nodes together. The red region corresponds to more informative for the score prediction. A, B) Visualizations training with all cortical and subcortical structures (All). C, D) Visualizations training with only cortical (Only-cortical) or subcortical structures (Only-subcortical).

**Fig. 4.**
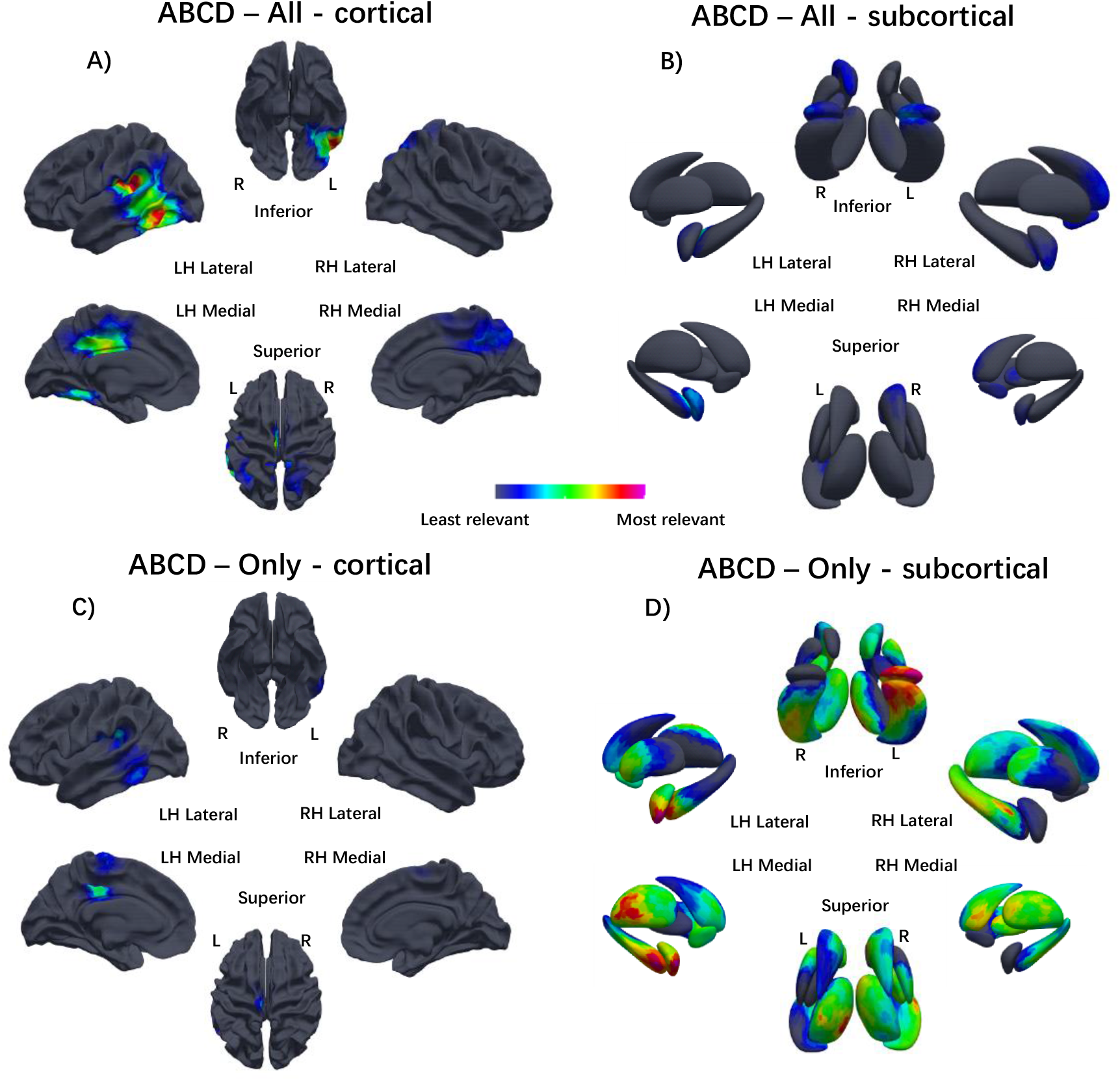
Grad-CAM visualizations on ABCD dataset using both cortical and subcortical structures as input. Brain regions predictve of fluid intelligence. Network trained on all cortical and subcortical nodes together. The red region corresponds to more informative for the score prediction. A, B) Visualizations training with all cortical and subcortical structures (All). C, D) Visualizations training with only cortical (Only-cortical) or subcortical structures (Only-subcortical).

Fig. 3A, B and Fig. 4A, B demonstrate that cortical structures play a more significant role than subcortical structures in the prediction of fluid intelligence score, which is in keeping with our statistical results. The topographic distribution of relevant brain structures is largely conserved with particular weight placed on the left temporal and parietal lobes in the prediction across both datasets. Interestingly, the morphology of the left temporal lobe was weighted more heavily in the prediction using the HCP dataset whereas the left parietal lobe was weighted more heavily in the prediction using the ABCD dataset. Other cortical structures including the bilateral paracentral lobules and posterior cingulate gyri were also relevant to the prediction but to a lesser degree. As the results in Table 1 and Table 2 demonstrate, there is more broadly distributed involvement of the subcortical structures in the prediction of fluid intelligence from the ABCD dataset while the subcortical structures are somewhat less involved in prediction from the HCP dataset.

Results from the models using only cortical surface data or only subcortical surface data were similar in distribution but variable in degree when compared to results from the model using both cortical and subcortical surface data together.

## Discussion

This study had three aims. These aims were to (a) identify the most predictive brain morphometric features of Gf, (b) assess the contribution of the subcortical morphometry to Gf either alone or combined with cortical morphology, and (c) investigate how age might be involved in the prediction of Gf. Our Analysis utilized a novel deep learning model using residual gCNNs to infer Gf from cortical and subcortical surface models that integrated multiple morphometric features such as cortical thickness, cortical area and gyrification, as well the shape of subcortical structures. Using two large and independent datasets of pre-adolescent (ABCD project) and young adults (HCP dataset) and a nested six-fold cross-validation scheme, this analysis predicted Gf with significant correlations (R=0.31-0.45). Across both datasets, the right amygdala and hippocampus, left temporal and parietal cortex, and bilateral cingulate morphometry consistently drove the prediction of Gf. Given the uniqueness of these findings, particularly for the amygdala and temporal cortex, localization was confirmed using grad-CAM to show reproducibility across subcortical surfaces and gyral folds. Divergence between the datasets was observed whereby the left hippocampus and amygdala, right caudate, right nucleus accumbens (NAc) and right pallidum also helped predict Gf for the younger ABCD cohort, with subcortical structures alone producing an R ~ 0.27, and cortical structures alone producing an R ~ 0.3. Together, subcortical and cortical structures produced an R ~ 0.31 for the younger ABCD cohort, whereas for the older HCP cohort, the combined was higher (R ~ 0.45). For the older HCP cohort, furthermore, subcortical structures alone producing an R ~ 0.1, and cortical structures alone produced an R ~ 0.4, with the right rectus gyrus helping predict Gf for the older HCP cohort but not the ABCD cohort. In both datasets, significantly better predictions were thus obtained by combining the cortical and subcortical surfaces. Despite this commonality regarding improved Gf prediction through combining cortical and subcortical morphometric features, the younger ABCD cohort had a substantially larger contribution from medial temporal and basal ganglia brain regions than the older HCP cohort, which needs to be considered in the context of differences in the trajectory of brain development between the late latency/pre-adolescence (ABCD cohort) and late adolescence/young adulthood (HCP cohort).

### Predictive models of Gf

Fluid intelligence refers to the ability to solve novel reasoning problems, which is considered independent of experience and education and, as such, believed to be biologically grounded on neurodevelopment (*40*). Previous studies have reported an age-related performance in Gf, peaking in late adolescence and declining in adulthood (*41*). In this study, we included two datasets of subjects at distinct phases of cognitive maturation. The younger cohort, the ABCD dataset, included children from 9 to 11 years, an age at which fluid intelligence has not yet reached its putative maximum. In this cohort, we predicted Gf with R= 0.328, which, to our knowledge, further improves the prediction accuracy so far reported using this dataset (*42–46*). Using Kernel Ridge Regression classifiers and CNNs, Mihalik and colleagues used manually extracted voxel-wise brain features (as opposed to automated morphometric analysis) on the ABCD dataset and predicted residualized Gf with an R = 0.17 (*43*), while Li and colleagues XGBoost classifiers on brain volumes and cortical curvatures to predict Gf with an R = 0.18 (*47*). The current work substantially builds on these ground-breaking reports, while also identifying brain region features, such as amygdala shape, not previously reported to be involved with Gf.

More studies have attempted to predict fluid intelligence using the HCP dataset. The age of subjects in the HCP dataset ranges from 22 to 35 years old, which corresponds to a different maturational phase when fluid intelligence is close to its full potential (*48*). All previous studies predicting fluid intelligence in the HCP dataset have done so using functional MRI (fMRI) (*49–52*). For instance, using functional connectivity analysis of task-based fMRI (FC), Greene and colleagues reached an R = 0.17 (*53*). Combining FC with resting-state fMRI (rs-fMRI), Elliott and colleagues obtained an R = 0.325 (*54*). Jiang and colleagues integrated multi-task FC features, applying partial least square regression method to improve the accuracy to an R = 0.409 (*55*). The current work compares favorably with these previously reported state-of-the-art functional imaging methods, by achieving an R = 0.454, using T1 weighted anatomic MRI data without any behavioral or functional imaging data. Our results support an association between brain morphometry and Gf (*21*). Moreover, we found that this association was strengthened when both cortical and subcortical structures’ shapes informed our gCNNs, underpinning the interdependencies across remote brain regions that in our review of the literature has not been reported.

### Cortical and subcortical regions involved in the prediction of Gf

The degree of involvement of temporal, parietal, and cingulate cortex, as revealed by grad-CAM, was highly reproducible across folds and displayed remarkable similarities between the two independent datasets. Specific cortical regions for both datasets included the left posterior middle and inferior temporal gyri as well as left basal temporal cortex, left temporo-parietal junction at the posterior aspect of the Sylvian fissure, left posterior cingulate, left interhemispheric paracentral lobule and the right cingulate region. At the cortical level, the only differentiator between the two datasets was the right rectus gyrus, whose morphometry predicted Gf in the HCP dataset but not in the ABCD dataset. These morphometric findings of temporal, parietal, and cingulate cortex add complexity to current frameworks for Gf that focus on fronto-parietal networks involved with combining sensory and executive material (*14*) as with parieto-frontal integration theory (P-FIT)(*15, 17*). The fact that morphometric features of the temporal, parietal, and cingulate cortex were observed in two independent cohorts raises many issues about integrating such structural imaging findings with functional imaging findings that focus on other regions of the prefrontal cortex, particularly as the current structural findings have as strong a predictive outcome as any study to date using functional imaging data, and question the current focus of many studies of Gf just on fronto-parietal networks.

Prior work of the neuroanatomic substrate of Gf has identified associations between widespread cortical areas, but relatively few relationships were reported with subcortical structures. The subcortical structure that appears to have had the most associations with Gf is the hippocampus. Raz and colleagues reported smaller hippocampal volume being associated with Gf (*56*) while Amat and colleagues reported smaller hippocampal volume being associated with full-scale intelligence quotient (IQ) and IQ subscales (*57*). Others reported hippocampal volume predicting Gf only in musically trained people (*58*), and the volumes of hippocampal subfields being more relevant for Gf than working memory (*59*), even though working memory has been linked to Gf (*11*). The current findings support the prior work, particularly in the context that increased Gf prediction resulted when subcortical regions such as the hippocampus were combined with cortical regions; this work resembles but does not replicate others who have reported that rs-fMRI connectivity between the right hippocampus and medial prefrontal cortex was associated with Gf (*60*). The current work further indicates how important it is to consider hippocampus morphometry in the context of the morphometry of other subcortical regions, particularly those with minimal association to Gf in the literature, that also have been linked to other cognitive science literatures, such as reward processing in judgement and decisionmaking and emotion regulation (e.g., nucleus accumbens and amygdala).

It appears a relatively smaller number of studies have linked Gf to morphometric measures of regions of the basal ganglia, such as the caudate and NAc (*22*), or suggested that Gf can be segregated from Gc based on NAc volume (*61*). The current work adds to these studies by indicating the bilateral NAc was important for predicting Gf in latency stage individuals (ABCD cohort), but contributes to the prediction of Gf to a lesser extent than cortex in late adolescents/young adults (HCP cohort). The NAc has been a fundamental target of addiction research (*62*), social reward studies (*63, 64*) and neuroeconomics (*65*), with a consensus sentiment that it is a core region for the judgement of value, that is fundamental for decision-making (*66, 67*). In this context, the NAc has also been considered important for allocation of effort, as with effortful cognitive tasks and motivation (*12*), and has been implicated in “grit” or the ability to persevere in a motivated fashion under adversity (*68*). The NAc is a critical target of dopaminergic cells in the brainstem (*67*), that make it important for motivated behavior, and suggest it would be important for allocating effort to the solution of novel reasoning problems that define Gf.

Related to the function of motivation, and heavily interconnected with the NAc (*12*) the amygdala has been considered a core region for emotion regulation, such as the experience and control of fear (*69*). To date, we cannot find any studies in the literature that implicate the amygdala with Gf, despite multiple studies implicating other regions with Gf that are contiguous with the amygdala (e.g., hippocampus) or significantly interconnected with it (e.g., NAc). Gf has been implicated with connectivity related to the uncinate fasciculus, a white matter bundle that connects the amygdala and anterior temporal cortex with frontal regions (*70*), but not directly connected to amygdala morphometry. The current findings across two independent cohorts of amygdala morphometry predicting Gf, might be consistent with a role in emotion regulation facilitating the solution of novel problems and adapting learning to new circumstances.

In parallel with considering the location of morphometric changes observed in this study, it is important to consider the complexity involved with morphometry as a field, including the number of independent features measured by voxel-based morphometry, cortical thickness, and volumetrics (*19, 23, 26, 27, 71, 72*). The analysis of the specific contributions of cortical thickness, cortical area and gyrification to Gf can reveal large topologic variations depending on the cortical morphometry employed, and resulting in sometimes contradictory results that suggest limitations to the specificity of each measurement individually (*21, 73, 74*). Using a data-driven approach which is agnostic to the individual morphologic features of the brain’s shape, the approach used in this study identified robust and well-localized involvement of both cortical and subcortical regions. Although the exact nature of the inferred morphometric features is not known using this approach, the network has the ability to identify interactions across individual morphologic features including cortical thickness, cortical area and gyrification, as well as to integrate features related to the shape of subcortical structures in its learning process. It can also take into account subtle and non-linear inter-regional interactions that contribute significantly to an individual’s Gf. Multiple brain regions previously reported in the literature using individual morphologic feature analysis were not revealed to play a role in the prediction of Gf using the current approach. One explanation for this is that our method integrates multi-dimensional interactions across individual morphologic features into its prediction, and the mapped results identified the most relevant brain regions taking these interactions into account.

### Differences in topographic prediction of Gf across age groups

Gf increases rapidly from birth through late adolescence, when it reaches a plateau which is sustained through the third decade of life, followed by a slow decay over the remaining lifespan (*75*). This trajectory parallels that of grey matter pruning in the cortex, which is much more pronounced in latency-aged children (e.g., ABCD cohort) relative to young adults (HCP cohort). Throughout adolescence, a strong relationship between cortical and subcortical development has been noted with cognitive performance (*76, 77*). Stress and emotional strain from adverse familial, educational, and social events over childhood and adolescence can also modulate the rate of growth in Gf (*78*). One might thus expect larger inter-subject variability in a younger population when Gf is still in its developmental phase rather than in a young adult population when its changes is asymptotic. Our results could be consistent with this interpretation in that we achieved a higher R in predicting Gf for late adolescents/young adults (HCP cohort) relative to latency stage/pre-adolescent children (ABCD cohort). At the same time, the cortical brain regions involved in the prediction of Gf remained consistent across age groups as revealed by grad-CAM visualization, despite the differences in predictive accuracy. Two other issues also should be noted. Namely, that neurodevelopment impacts the capacity to modulate cognitive behaviors important for Gf (*79, 80*). Furthermore, subjects from the HCP dataset were all healthy adults while the ABCD dataset included children with a broad array of risk factors for developing mental health and addictive disorders, which can impact Gf (*81, 82*) Differences in the discrepancy in accuracy across datasets likely represents contributions from a combination of the brain’s developmental trajectory as well as potential cognitive vulnerabilities across the health spectrum.

Between these two cohorts, subcortical structures played a more prominent role in the prediction of Gf in latency-stage/pre-adolescent children than in young adults/late adolescents. Across both cohorts, only the head of the right hippocampus and the right amygdala consistently contributed to the prediction of Gf. For the younger subjects (ABCD cohort), the left hippocampus and amygdala were also important for the prediction of Gf, along with the right caudate, NAc, and pallidum. The observation of bilateral hippocampi with the ABCD cohort is consistent with suggestions that working memory may be particularly important for Gf in children (*83*). In the developing brain, associations between fluid reasoning and subcortical shape has been reported to be widespread, encompassing the bilateral putamen, pallidum and caudate (*84*), consistent with the current findings. Our findings involving the right striatum are in keeping with other reports of asymmetric right-sided striatal dominance in younger individuals compared to older individuals (*85*). Lastly, it needs to be noted that medial temporal structures and the striatum have strong connections to frontal and cingulate cortices (*86*), and corticostriatal circuits (*87*). Through such connections medial temporal structures and the striatum have been implicated with executive function (*88*), context coding (*89*) and impulse control (*90*), which are important processes for adaptation to novelty with Gf.

## Materials and Methods

### Datasets

Brain MRI and neurocognitive data from two publicly available datasets were used independently in this work: the Human Connectome Project (HCP) S1200 data release and the Adolescent Brain Cognitive Development Study (ABCD) 2.0 release (*82, 91*). The HCP dataset consists of neurobehavioral measurements and MRI scans from healthy subjects aged between 22 to 35 years. Subjects were defined as “healthy” in the absence of diagnosed neurologic or psychological conditions. All subjects were scanned on a custom Siemens 3T Connectome Skyra at Washington University using a standard 32-channel Siemens head coil. Further details pertaining to the included subjects, data collection parameters and preprocessing steps can be found on the HCP website (*82, 92*). The ABCD dataset consists of neurobehavioral measurements and MRI scans from over 11,000 children aged between 9 and 11 years. Subjects from across the United States with diverse health, socioeconomic and ethnic backgrounds were included. Brain MRI data were acquired from three different 3T scanner platforms: Siemens Prisma, General Electric 750 and Phillips. Further details pertaining to the included subjects, data collection parameters and preprocessing steps can be found on the ABCD website (*91*). Minimally preprocessed T1-weighted MRI scans were obtained from both databases.

In addition to brain MRI data, Gf scores, measured by the NIH Toolbox Neurocognition battery were collected. Specifically, the *nihtbx_fluidcompuncorrected* variable was included from the ABCD dataset and the *CogFluidComp_Unadj* variable was included from the HCP dataset (*93*). This Toolbox Fluid Cognition Composite score was computed by the average of the raw scores from six measures of fluid abilities (the Toolbox Dimensional Change Card Sort [DCCS] Test, the Toolbox Flanker Inhibitory Control and Attention Test, the Toolbox Picture Sequence Memory Test, the Toolbox List Sorting Working Memory Test, and the Toolbox Pattern Comparison Processing Speed Test). The raw Gf scores from two datasets were quantile normalized at first in order to assume the Gaussian distribution of each dataset. Quantile normalization was realized by sorting the scores of each subject from low to high and replacing them with a random standard Gaussian distribution which was also sorted from low to high. The characteristics of two datasets are summarized in Table 3.

**Table 3.** The characteristics of HCP and ABCD datasets.

|  | **HCP**<br>**(N = 1097)** | **ABCD**<br>**(N = 8070)** |
| --- | --- | --- |
| **Age**<br>(mean ± Sd <sup>a</sup>) | 28.81±3.70 | 9.93±0.62 |
| **Sex**<br>(Female/male) | 596/501 | 3861/4209 |
| **Fluid intelligence**<br>(mean ± Sd) | 115.07±11.58 | 92.25±10.43 |
| **Health status** | All good | Different conditions |
| **Scanner** | Siemens 3T<br>Connectome Skyra | Siemens Prisma, General<br>Electric 750 and Phillips |

<sup>a</sup>. Sd: Standard deviation.

### MRI Data Preprocessing

For each subject, inner cortical surfaces (modeling the interface between grey and white matter) and outer cortical surfaces (modeling the cerebrospinal fluid/grey matter interface) were extracted using Freesurfer (v6.0, https://surfer.nmr.mgh.harvard.edu). Seven subcortical structures per hemisphere were automatically segmented using Freesurfer (amygdala, nucleus accumbens, caudate, hippocampus, pallidum, putamen, thalamus) and then modeled into surface meshes using SPHARM-PDM (https://www.nitrc.org/projects/spharm-pdm/). All surfaces were inflated, parameterized and registered to a corresponding surface template using a rigid-body registration to preserve the anatomy of the cortex and subcortical structures (*94*). No morphometric evaluation of subcortical structures, re-segmentation, or use of multiple atlases was performed; this study sought to minimize variance from analysis of the feature set used for prediction.

Surface templates were converted to graphs based on their triangulation scheme. Nodes of the graphs were surface vertices, and edges of the graphs were segments across vertices. Overall, the graphs including all structures had 47,616 nodes, 32,768 from the cortical surfaces and 14,848 from the subcortical surfaces. Input features of the network were defined as the Cartesian coordinates of surface vertices in subjects’ native space resampled into the surface templates. As a consequence, cortical nodes were assigned 6 features (X, Y, Z of both the inner and outer cortical surface vertices) and subcortical nodes had 3 features (X, Y, Z of subcortical surface vertices) when they were used for separate training.

More details about the construction of the common graphs and the organization of input features are provided in Supplemental material and Fig. S1–4. All subjects were represented using the same underlying graphs, the features assigned to the nodes were unique to each subject and were the input of our gCNNs.

### Preparation of the datasets

The nested cross-validation was used in this work to assess generalizability, which contains an outer loop of six folds and an inner loop of five folds. Both datasets were split into six folds, randomly selecting one set as the outer test set and the rest five sets as the outer training set. This whole process repeats six times for each fold. The outer training set was sub-divided into five folds, including one validation set and four inner training sets. This inner process repeated five times and the outer test set was evaluated by an ensembled model averaged from those five trained models. More details are shown in Fig. S2. For HCP dataset, we included 1,097 subjects, i.e., in each fold, 914 inner training sets and 183 outer test sets and for ABCD dataset, we included 8,070 subjects, i.e., 6,725 inner training sets and 1,345 outer test set.

### Graph Convolution

Traditional CNNs extract features on structured data, such as 2D images or 3D volumes. The convolution operations over graphs can be generalized in the spectral domain, which is the multiplication of the signal on graphs with the eigenvector matrix of the graph Laplacian (*95*). We defined un undirected graph *G* = {*V, ϵ, A*}, where edge ϵ connects two vertices V, with |*V*| = *n*, and *A* ∈ *R^n ×n^* is the weighted adjacency matrix. *A* is a square symmetric matrix and *A_ij_* is the weight assigned to edge (*i,j*) that connects vertices *i* and *j*. Then the graph Laplacian is defined as *L* = *D* – *A* and its normalized form is given by 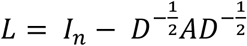, where 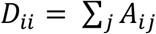 is an entry of the diagonal degree matrix D of the graph and *I_n_* is the identity matrix. *L* is a diagonalizable matrix that can be factored using the eigen-decomposition *L* = *U*Λ*U^T^*, where *Λ* = *diag* ([*λ*_0_, *λ*_1_,…, *λ*_*n*–1_]) ∈ *R^n×n^* is the diagonal matrix of eigenvalues and *U* = [*u*_0_, *u*_1_,…, *u*_*n*–1_] ∈ *R^n×n^* is the matrix containing the set of corresponding orthogonal eigenvectors.

The graph Fourier Transform (GFT) of our input feature matrix *X* ∈ *R^n ×f^* is defined as 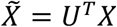 and its inverse as 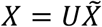, where *f* = 3,6 *or* 9 is the number of features. Given a filter *g_θ_* = *diag*(*θ*) and an arbitrary graph signal *x*, let us define by *y* = *x* * *g* the convolution between *x* and *g*. This corresponds to multiplication in the Fourier space, that is,

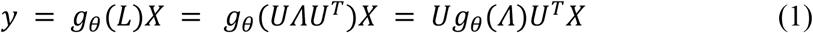

where *θ* ∈ *R^n^* is a vector of Fourier coefficient.

In order to largely reduce the computational complexity, we approximate spectral filters using truncated expansions of Chebyshev polynomials (*95*). The *K*-localized filtering operation is defined as

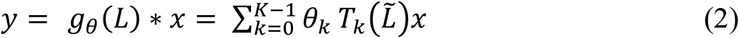

with scaled 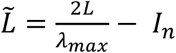, where *λ_max_* denotes the largest eigenvalue of *Λ* and *θ_k_* represents the *k*-th Chebyshev coefficient. 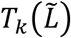 is the Chebyshev polynomial of order *k*, which is calculated by 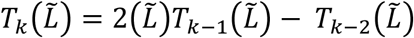, where 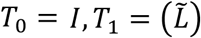. Then the number of trainable parameter per layer is reduced to *F_in_* × *F_out_* × *K*, where *F_in_, F_out_* are the number of corresponding input and output features.

### Image Augmentation

In order to increase the generalizability between our two datasets, we used two data augmentation techniques: random rotation within ±20 degrees and a random Gaussian noise standardized with mean *μ* = 0 and a standard deviation of *σ* = 0.02. The augmentation parameter *p_a_* denotes the probability of the use of data augmentation for a single subject. In this study, both datasets had *p_a_* = 0.5, indicating that data augmentation was applied with a 50% probability per subject for each iteration of training.

### Network Architecture

Fig. 5 shows the details of our graph networks. We used residual blocks in our model to facilitate the training of deeper networks inspired by (*96*). Using this approach, the output of the previous block is added to the output of the current block to avoid the vanishing gradient problem (*97*). Our model contains a pre-convolutional layer (Pre-Conv), four residual blocks (ResBlock) and a post residual block, followed by a single fully connected (Fc) layer with one output that reflects the estimated Gf score. Each residual block has two subblocks, including a batch normalization layer (BN), a non-linear activation function ReLU and a normal convolutional layer (Conv). The maxpooling layer is used after each residual block to downsample the features.

**Fig. 5.**
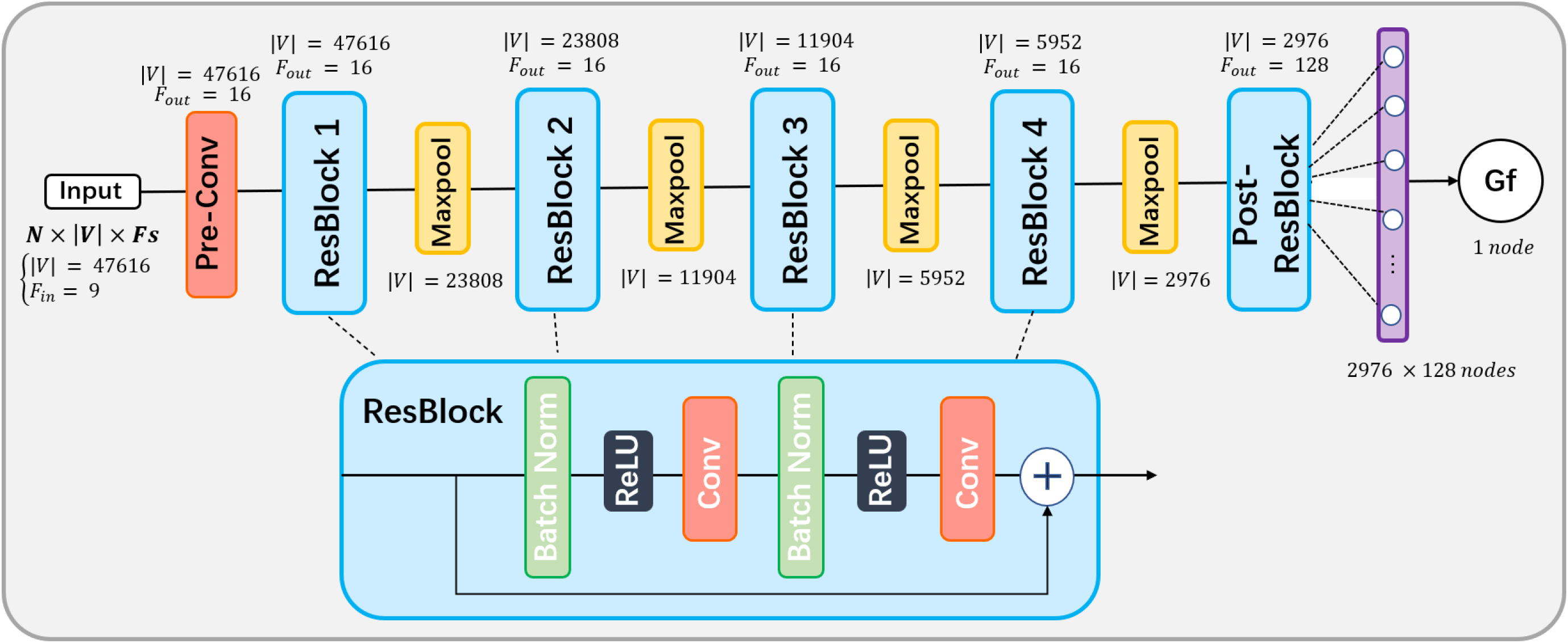
The model architecture. The model contains a pre-convolutional layer, four residual blocks and a post residual block, followed by a fully connected layer; Each residual block has two subblocks, each with a batch normalization layer, a ReLU and a convolutional layer. Each residual block is followed by a maxpooling layer to downsample the features. N is the number of batch size; |*V*| is the number of vertices; F is the number of features.

### Loss function

The loss function used in our model is composed of three parts: the mean square error (MSE) between the network’s estimations and actual values, the Pearson’s coefficient of correlation *corr* and additional *l*_2_ regularization term. *L_all_* is defined as

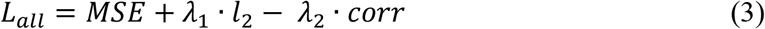

for 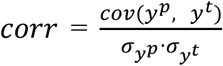, where *λ*_1_, *λ*_2_ are the regularization parameters, *y^p^* is the predicted score, *y^t^* is the true label, *cov* is the covariance and *σ* is the standard deviation. This correlation term is added to alleviate the “regression towards or to the mean (RTM)” bias, where the higher the correlation, the lower the loss (*98*).

### Grad-CAM Visualization

To visualize the most relevant brain areas involved in the network’s decision making process and to provide some interpretability to our network results, a graphical gradient-weighted Class Activation Map (grad-CAM) method was applied to generate a color-coded probability map *M_c_* (*99*). Grad-CAM uses the gradient information flowing back to the last convolutional layer of the model to generate heatmaps highlighting important regions upon which the model focuses and then performs a global average pooling operation to produce the importance weights 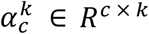, of each neuron:

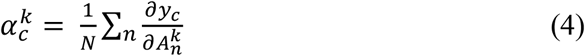

where *y_c_* refers to the score of class *c*, i.e. *c* = 1 in this paper, and 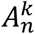 represents the value at each node *n* for feature map activations *A^k^* on the last convolutional layer. After getting the weights, *M_c_* is calculated with a weighted combination of feature maps followed by an activation function *ReLU*, which is applied to only take positive weights into consideration and ignore the negative weights since we were interested in the features with positive influence on the class of interest.

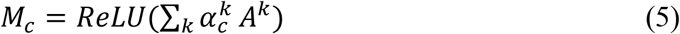

Grad-CAM maps were obtained for Gf prediction from each testing set in all six folds. In our models we used four pooling layers, reducing the number of nodes by a factor 2^4^. Therefore, we rescaled the generated grad-CAM maps back to the original size using spherical linear interpolation on the cortical and subcortical surfaces in order to overlay on the original graphs.

### Network implementation

Model performance was evaluated using nested cross-validation, with the datasets split into six folds each, where each fold was randomly chosen for testing and the remaining five folds were used for training. In each fold, the outer test dataset was evaluated by an averaged model ensembled from all five inner-folds models, and the grad-CAMs were generated using the average weighted sum on each of the testing subject. More details are shown in Fig. S2.

For the ABCD dataset, the networks were trained using a batch size of 32, and a maximum number of epochs of 100. We used Adam optimizer with a learning rate of 0.0005 and a decay rate of 0.99 per ten steps, the parameters *λ*_1_ and *λ*_2_ were both set to 0.0001 and the dropout ratio at the fully connected layer was set to 0.5. For the HCP dataset, the batch size was set to 50 and the parameter *λ*_1_ was set to 0.0005. Due to the smaller dataset size, the maximum number of epochs for the HCP dataset was set to 80. The different network parameters were optimized using cross-validation and the training process stopped when the generalization error increased with the patience factor set to 5. The networks were implemented in Python (version 3.6) using TensorFlow and trained on a single GPU (Nvidia GeForce 2080Ti) workstation.

### Statistical Analysis

The performance of three types of gCNNs were evaluated, using either: 1) only the inner and outer cortical surface nodes, 2) only the subcortical surface nodes, or 3) both inner and outer cortical surface and subcortical surface nodes together. The MSE, Pearson correlation coefficient score (R) and training time yielded for each testing fold and for each complete dataset were calculated. A paired t-test was performed to compare the performance of each of the three input types and the p-values were adjusted for multiple comparisons using false discovery rate (FDR), which was considered as statistically significant if the p-values < 0.05. The normalized correlation (0 to 1) was calculated on the mapping results (*M_c_*) to compare the inter-fold similarity and inter-cohort similarity on the HCP and ABCD datasets. The closer to 1, the higher the correlation between the two groups. All statistical analysis was performed using the Scikit-learn and Numpy packages in Python (version 3.6).

## Acknowledgments

This research was supported in part through the computational resources and staff contributions provided for the Quest high performance computing facility at Northwestern University which is jointly supported by the Office of the Provost, the Office for Research, and Northwestern University Information Technology.

## Competing interests

The authors declare no competing interests.

## Supplementary Materials

### Materials and Methods

#### Conversion of meshes to graphs

Template surface meshes were converted to graphs using their triangulation schemes (Fig. S3). Nodes of the graphs were defined as the surface vertices, and edges of the graphs were triangles segments across vertices. A weight *W_i,j_* was assigned to the edge between neighbor nodes *i* and *j* such as:

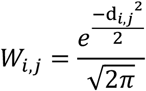

with *d_i,j_* is the Euclidean distance between the nodes *i* and *j*.

#### Hierarchical decomposition of the graphs

High-resolution template graphs underwent a hierarchical dichotomic partitioning using the following steps and illustrated in Fig. S4:

1. For each structure, define the root leaf as the graph corresponding to the high-density mesh
2. Repeat these steps until the average distance across neighbor leaves is less than a threshold *T*

a. Given a set of leaves at level *L*, partition each leaf at level *L* into two child leaves at level *L* + 1 using spectral clustering (*100*). The new leaves are therefore subgraphs of their parent leaf;
b. Order the 2^*L*+1^ leaves of level *L* + 1 such that the leaves 2(*i* – 1) and 2*i* are partitions of leaf *i* at level *L*;
c. For each leaf of level *L* + 1, identify its center node as the node whose betweenness centrality is largest (*101*);
d. Defines the partition neighbor matrix *M*_*L*+1_, of size 2^*L*+1^ × 2^*L*+1^, such that

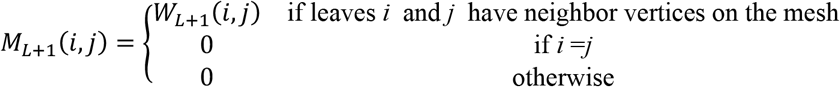

and

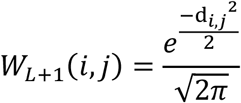

where *d_i,j_* is the geodesic distance (along the mesh) between the center of the leaf *i* and the center of the leaf *j* at level *L* + 1;
e. Compute the average distance across leaves’ centers and exit the loop if less than *T*, otherwise continue to step a.

The average distance threshold *T* was set to 3 mm for the cortical surfaces and 2 mm for the subcortical surfaces. Therefore, the number of decomposition levels for each structure was a function of their surface area. The number of nodes at the finest level is provided in Table S2.

#### Aggregation of matrices and features

Depending on the structures used for the prediction, the matrices *M_L_* were block concatenated along the main diagonal to define the whole underlying graph (Fig. S5).

When only the subcortical structures were used for prediction, the input features associated to the nodes of the graph were the 3-dimensional Cartesian coordinates of the corresponding center node in individuals’ space [*X_s_, Y_S_, Z_S_*]. If the cortex only was used to feed the gCNN, then a 6-dimensional vector was assigned to each node of the graph, containing the Cartesian coordinates of the inner and outer cortical surfaces [*X_w_, Y_w_, Z_w_, X_p_, Y_p_, Z_p_*]. Finally, when both the subcortical structures and the cortex were used to feed the gCNN, a 9-dimensional vector was assigned to each node of the graph. For nodes of the subcortical structures, the first 3 elements of the input were Cartesian coordinates of the nodes, and the last 6 were zeros. For cortical nodes, the first 3 elements of the vector were zeros, and the last 6 the Cartesian coordinates of the inner and outer cortical surfaces.

#### Pooling operation

By construction of the hierarchical decomposition of the graph and ordering of the node, the pooling operator is applied similarly to a 1-dimensional signal with a stride of 2 and a pooling size of 2. This is conceptually identical to the original gCNN study (*95*). However, as opposed to the METIS algorithm initially proposed, our approach guaranties that no singleton is ever generated.

**Fig. S1.**
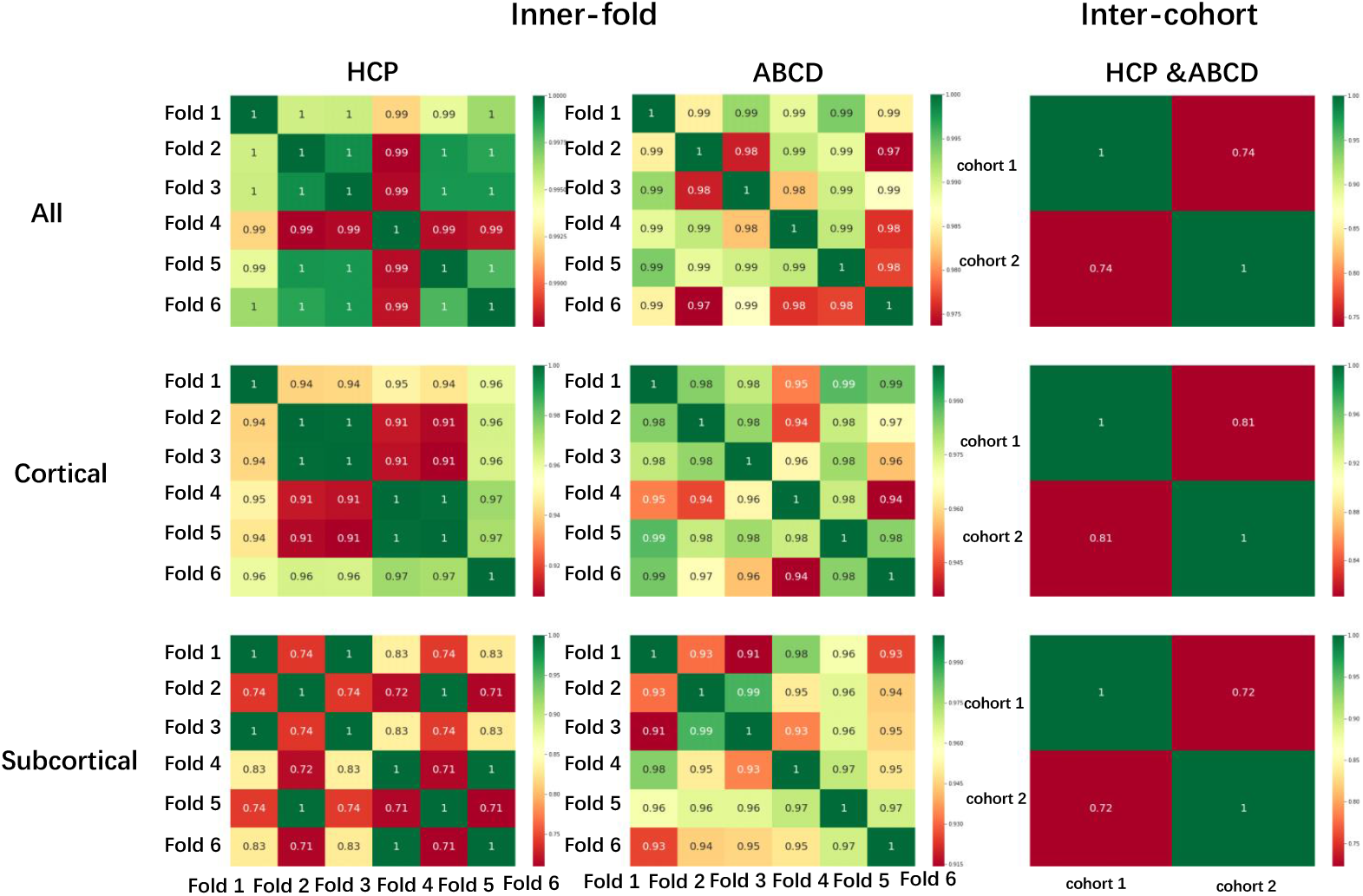
The inner-fold and inter-cohort correlation details of mappings (*M_c_*) on HCP and ABCD datasets. The closer to 1, the higher correlations of two groups.

**Fig. S2.**
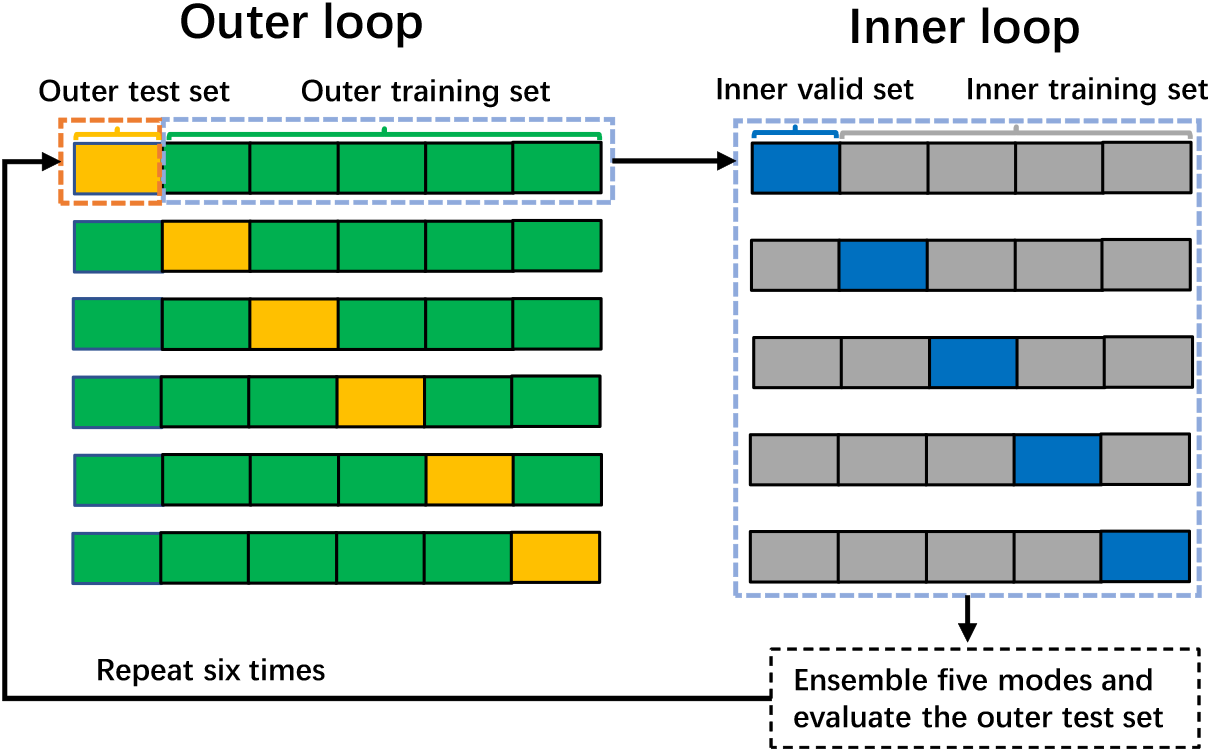
Illustration the training and evaluation process in six-fold nested cross-validation. The whole process contains an outer loop of six folds and an inner loop of five folds. The model is trained on inner training sets, finetuned on inner validation sets and evaluated on the outer test sets.

**Fig. S3.**
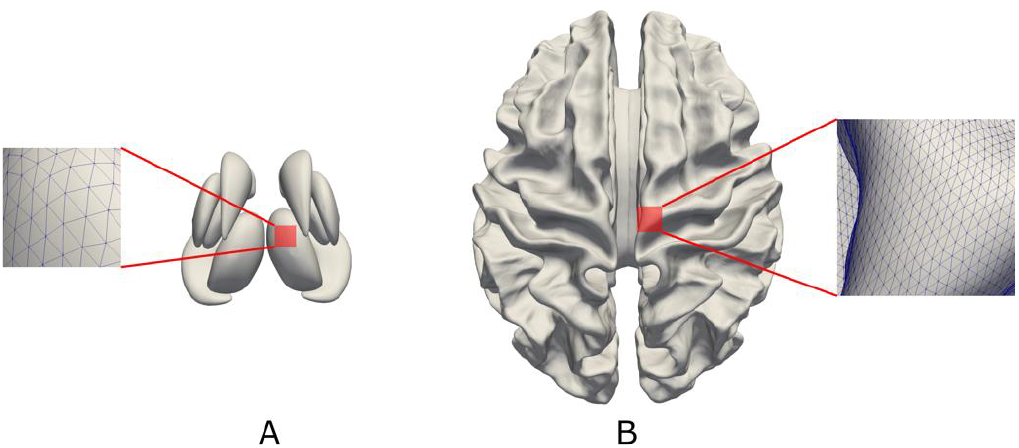
Triangulation schemas of surface meshes. Visualization of the surface template for the subcortical structures (A) and the cortex (B), with close up on their high-density triangulations.

**Fig. S4.**
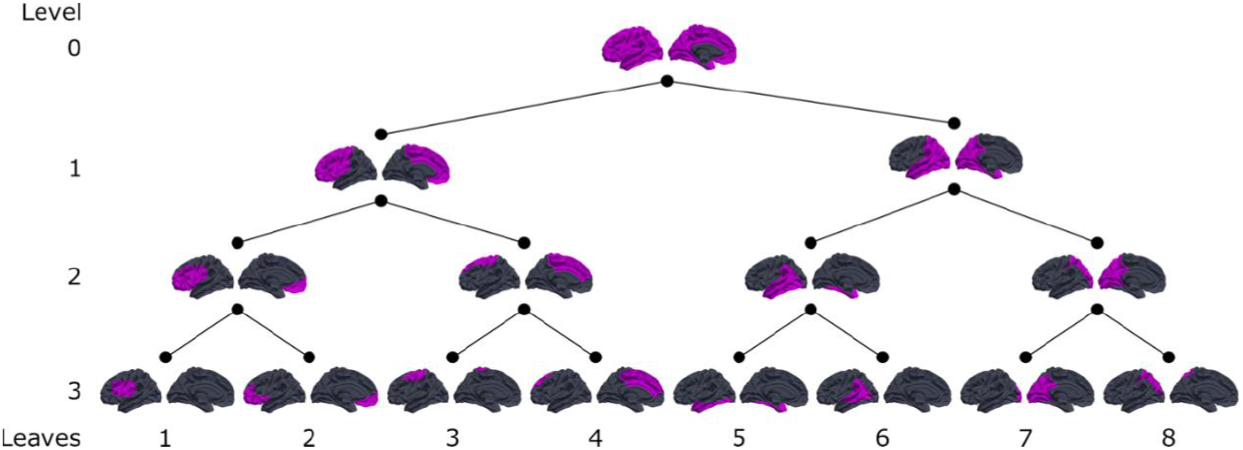
First four level of the hierarchical decomposition of the left cortical surface. The initial level (0) is the whole structure, in that case the left cortical surface masking out non-cortical regions such as Freesurfer’s medial wall and parahypocampal regions.

**Fig. S5.**
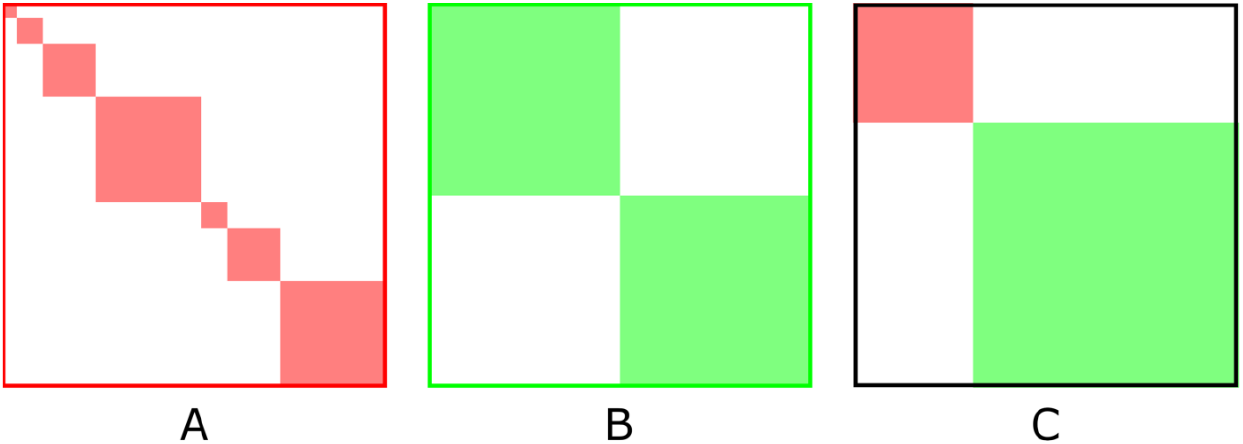
Block concatenation of individual structures matrices *M_L_* along the main diagonal. A) Concatenation of the seven subcortical structures matrices along the main diagonal used for fluid intelligence prediction with the subcortical structures only; B) Concatenation of the left and right cortical hemisphere matrices along the main diagonal for the fIQ prediction with the cortex only; C) Concatenation of the subcortical (A) and cortical (B) matrices for the prediction using all the structures

**Table S1.** The averaged correlations of mappings (*M_c_*) on HCP and ABCD datasets.

|  | Inner- fold |  | Inter- cohort |
| --- | --- | --- | --- |
|  | HCP | ABCD | HCP & ABCD |
| All <sup>a</sup> | 0.996 | 0.988 | 0.742 |
| Cor <sup>b</sup> | 0.957 | 0.975 | 0.814 |
| Sub <sup>c</sup> | 0.841 | 0.960 | 0.721 |

<sup>a</sup>. All: use both cortical and subcortical nodes. <sup>b</sup>. Cor: cortical nodes only. <sup>c</sup>. Sub: subcortical nodes only

**Table S2.** Number of nodes for each structure at their finest decomposition level.

| Structure | Number of nodes<br>Per hemisphere |
| --- | --- |
| Accumbens | 256 |
| Amygdala | 512 |
| Caudate | 1024 |
| Hippocampus | 2048 |
| Pallidum | 512 |
| Putamen | 1024 |
| Thalamus | 2048 |
| Cortex | 16384 |

## Notes

### Competing Interest Statement

The authors have declared no competing interest.

## References

1. A. Binet, Th. Simon, A method of measuring the development of intelligence of young children (Chicago Medical Book Company, Chicago, IL, US, 1915), A method of measuring the development of intelligence of young children.

2. R. Siegler, The other Alfred Binet. Dev. Psychol. 28, 179–190 (1992).

3. L. S. Gottfredson, Why g matters: The complexity of everyday life. Intelligence. 24, 79–132 (1997).

4. R. J. Sternberg, Wisdom, intelligence, and creativity synthesized (Cambridge University Press, New York, NY, US, 2003), Wisdom, intelligence, and creativity synthesized.

5. R. B. Cattell, Theory of fluid and crystallized intelligence: A critical experiment. J. Educ. Psychol. 54, 1–22 (1963).

6. J. K. Hartshorne, L. T. Germine, When Does Cognitive Functioning Peak? The Asynchronous Rise and Fall of Different Cognitive Abilities Across the Life Span. Psychol. Sci. 26, 433–443 (2015).

7. J. L. Horn, G. Donaldson, R. Engstrom, Apprehension, Memory, and Fluid Intelligence Decline in Adulthood. Res. Aging. 3, 33–84 (1981).

8. T. A. Salthouse, What and When of Cognitive Aging. Curr. Dir. Psychol. Sci. 13, 140–144 (2004).

9. W. R. Cunningham, V. Clayton, W. Overton, Fluid and Crystallized Intelligence in Youni Adulthood and Old Age, 3.

10. A. F. Fry, S. Hale, Relationships among processing speed, working memory, and fluid intelligence in children. Biol. Psychol. 54, 1–34 (2000).

11. I. Deary, Why do intelligent people live longer? Nature. 456, 175–176 (2008).

12. H. C. Breiter, B. R. Rosen, Functional Magnetic Resonance Imaging of Brain Reward Circuitry in the Human. Ann. N. Y. Acad. Sci. 877, 523–547 (1999).

13. G. P. Gasic, J. W. Smoller, R. H. Perlis, M. Sun, S. Lee, B. W. Kim, M. J. Lee, D. J. Holt, A. J. Blood, N. Makris, D. K. Kennedy, R. D. Hoge, J. Calhoun, M. Fava, J. F. Gusella, H. C. Breiter, BDNF, relative preference, and reward circuitry responses to emotional communication. Am. J. Med. Genet. B Neuropsychiatr: Genet. 150B, 762–781 (2009).

14. E. Santarnecchi, A. Emmendorfer, S. Tadayon, S. Rossi, A. Rossi, A. Pascual-Leone, Network connectivity correlates of variability in fluid intelligence performance. Intelligence. 65, 35–47 (2017).

15. R. E. Jung, R. J. Haier, The Parieto-Frontal Integration Theory (P-FIT) of intelligence: Converging neuroimaging evidence. Behav. Brain Sci. 30, 135–154 (2007).

16. A. Woolgar, J. Duncan, F. Manes, E. Fedorenko, Fluid intelligence is supported by the multiple-demand system not the language system. Nat. Hum. Behav. 2, 200–204 (2018).

17. J. Duncan, The multiple-demand (MD) system of the primate brain: mental programs for intelligent behaviour. Trends Cogn. Sci. 14, 172–179 (2010).

18. G. Nave, W. H. Jung, R. Karlsson Linnér, J. W. Kable, P. D. Koellinger, Are Bigger Brains Smarter? Evidence From a Large-Scale Preregistered Study. Psychol. Sci. 30, 43–54 (2019).

19. Q.-Y. Gong, V. Sluming, A. Mayes, S. Keller, T. Barrick, E. Cezayirli, N. Roberts, Voxel-based morphometry and stereology provide convergent evidence of the importance of medial prefrontal cortex for fluid intelligence in healthy adults. NeuroImage. 25, 1175–1186 (2005).

20. S. Karama, R. Colom, W. Johnson, I. J. Deary, R. Haier, D. P. Waber, C. Lepage, H. Ganjavi, R. Jung, A. C. Evans, Cortical thickness correlates of specific cognitive performance accounted for by the general factor of intelligence in healthy children aged 6 to 18. NeuroImage. 55, 1443–1453 (2011).

21. E. Tadayon, A. Pascual-Leone, E. Santarnecchi, Differential Contribution of Cortical Thickness, Surface Area, and Gyrification to Fluid and Crystallized Intelligence. Cereb. Cortex. 30, 215–225 (2020).

22. M. Burgaleta, P. A. MacDonald, K. Martínez, F. J. Román, J. Álvarez-Linera, A. R. González, S. Karama, R. Colom, Subcortical regional morphology correlates with fluid and spatial intelligence. Hum. Brain Mapp. 35, 1957–1968 (2014).

23. I. Wartenburger, E. Kühn, U. Sassenberg, M. Foth, E. A. Franz, E. van der Meer, On the relationship between fluid intelligence, gesture production, and brain structure. Intelligence. 38, 193–201 (2010).

24. T. Heimann, H.-P. Meinzer, Statistical shape models for 3D medical image segmentation: A review. Med. Image Anal. 13, 543–563 (2009).

25. J. G. Csernansky, M. K. Schindler, N. R. Splinter, L. Wang, M. Gado, L. D. Selemon, D. Rastogi-Cruz, J. A. Posener, P. A. Thompson, M. I. Miller, Abnormalities of Thalamic Volume and Shape in Schizophrenia. Am. J. Psychiatry. 161, 896–902 (2004).

26. N. Makris, M. Oscar-Berman, S. K. Jaffin, S. M. Hodge, D. N. Kennedy, V. S. Caviness, K. Marinkovic, H. C. Breiter, G. P. Gasic, G. J. Harris, Decreased Volume of the Brain Reward System in Alcoholism. Biol. Psychiatry. 64, 192–202 (2008).

27. J. M. Gilman, J. K. Kuster, S. Lee, M. J. Lee, B. W. Kim, N. Makris, A. van der Kouwe, A. J. Blood, H. C. Breiter, Cannabis Use Is Quantitatively Associated with Nucleus Accumbens and Amygdala Abnormalities in Young Adult Recreational Users. J. Neurosci. 34, 5529–5538 (2014).

28. D. M. Coscia, K. L. Narr, D. G. Robinson, L. S. Hamilton, S. Sevy, K. E. Burdick, H. Gunduz-Bruce, J. McCormack, R. M. Bilder, P. R. Szeszko, Volumetric and shape analysis of the thalamus in first-episode schizophrenia. Hum. Brain Mapp. 30, 1236–1245 (2009).

29. M. P. Harms, L. Wang, D. Mamah, D. M. Barch, P. A. Thompson, J. G. Csernansky, Thalamic Shape Abnormalities in Individuals with Schizophrenia and Their Nonpsychotic Siblings. J. Neurosci. 27, 13835–13842 (2007).

30. M. P. Harms, L. Wang, C. Campanella, K. Aldridge, A. J. Moffitt, J. Kuelper, J. T. Ratnanather, M. I. Miller, D. M. Barch, J. G. Csernansky, Structural abnormalities in gyri of the prefrontal cortex in individuals with schizophrenia and their unaffected siblings. Br. J. Psychiatry. 196, 150–157 (2010).

31. D.-H. Kang, S. H. Kim, C.-W. Kim, J.-S. Choi, J. H. Jang, M. H. Jung, J.-M. Lee, S. I. Kim, J. S. Kwon, Thalamus surface shape deformity in obsessive-compulsive disorder and schizophrenia. NeuroReport. 19, 609–613 (2008).

32. M. J. McKeown, A. Uthama, R. Abugharbieh, S. Palmer, M. Lewis, X. Huang, Shape (but not volume) changes in the thalami in Parkinson disease. BMC Neurol. 8, 8 (2008).

33. L. Wang, D. Y. Lee, E. Bailey, J. M. Hartlein, M. H. Gado, M. I. Miller, K. J. Black, Validity of large-deformation high dimensional brain mapping of the basal ganglia in adults with Tourette syndrome. Psychiatry Res. Neuroimaging. 154, 181–190 (2007).

34. N. Makris, G. P. Gasic, D. N. Kennedy, S. M. Hodge, J. R. Kaiser, M. J. Lee, B. W. Kim, A. J. Blood, A. E. Evins, L. J. Seidman, D. V. Iosifescu, S. Lee, C. Baxter, R. H. Perlis, J. W. Smoller, M. Fava, H. C. Breiter, Cortical Thickness Abnormalities in Cocaine Addiction—A Reflection of Both Drug Use and a Pre-existing Disposition to Drug Abuse? Neuron. 60, 174–188 (2008).

35. V. B. Mountcastle, Perceptual Neuroscience: The Cerebral Cortex (Harvard University Press, 1998).

36. J. W. Prothero, J. W. Sundsten, Folding of the Cerebral Cortex in Mammals. Brain. Behav. Evol. 24, 152–167 (1984).

37. E. G. Jones, in Comparative Structure and Evolution of Cerebral Cortex, Part I, E. G. Jones, A. Peters, Eds. (Springer US, Boston, MA, 1990; https://doi.org/10.1007/978-1-4757-9622-3_9), Cerebral Cortex, pp. 311–362.

38. A. J. Rockel, R. W. Hiorns, T. P. S. Powell, The Basic Uniformity in Structure of the Neocortex. Brain. 103, 221–244 (1980).

39. P. Besson, T. Parrish, A. K. Katsaggelos, S. K. Bandt, “Geometric deep learning on brain shape predicts sex and age” (preprint, Neuroscience, 2020), doi:10.1101/2020.06.29.177543.

40. C. Blair, How similar are fluid cognition and general intelligence? A developmental neuroscience perspective on fluid cognition as an aspect of human cognitive ability. Behav. Brain Sci. 29, 109–125 (2006).

41. J. M. Bugg, N. A. Zook, E. L. DeLosh, D. B. Davalos, H. P. Davis, Age differences in fluid intelligence: Contributions of general slowing and frontal decline. Brain Cogn. 62, 9–16 (2006).

42. J. Li, R. Kong, R. Liégeois, C. Orban, Y. Tan, N. Sun, A. J. Holmes, M. R. Sabuncu, T. Ge, B. T. T. Yeo, Global signal regression strengthens association between resting-state functional connectivity and behavior. NeuroImage. 196, 126–141 (2019).

43. A. Mihalik, M. Brudfors, M. Robu, F. S. Ferreira, H. Lin, A. Rau, T. Wu, S. B. Blumberg, B. Kanber, M. Tariq, M. D. M. E. Garcia, C. Zor, D. I. Nikitichev, J. Mourao-Miranda, N. P. Oxtoby, ABCD Neurocognitive Prediction Challenge 2019: Predicting individual fluid intelligence scores from structural MRI using probabilistic segmentation and kernel ridge regression. ArXiv190510831 Q-Bio Stat (2019) (available at http://arxiv.org/abs/1905.10831).

44. N. P. Oxtoby, F. S. Ferreira, A. Mihalik, T. Wu, M. Brudfors, H. Lin, A. Rau, S. B. Blumberg, M. Robu, C. Zor, M. Tariq, M. D. M. E. Garcia, B. Kanber, D. I. Nikitichev, J. Mourao-Miranda, ABCD Neurocognitive Prediction Challenge 2019: Predicting individual residual fluid intelligence scores from cortical grey matter morphology. ArXiv190510834 Q-Bio Stat (2019) (available at http://arxiv.org/abs/1905.10834).

45. A. Wlaszczyk, A. Kaminska, A. Pietraszek, J. Dabrowski, M. A. Pawlak, H. Nowicka, in Adolescent Brain Cognitive Development Neurocognitive Prediction, K. M. Pohl, W. K. Thompson, E. Adeli, M. G. Linguraru, Eds. (Springer International Publishing, Cham, 2019), Lecture Notes in Computer Science, pp. 83–91.

46. Q. Zhao, L. Zhang, C. Shen, W. Cheng, J. Feng, Association Between Intelligence and Cortical Thickness in Adolescents: Evidence from the ABCD Study, 11.

47. T. Li, X. Wang, T. Luo, Y Yang, B. Zhao, L. Yang, Z. Zhu, H. Zhu, in Adolescent Brain Cognitive Development Neurocognitive Prediction, K. M. Pohl, W. K. Thompson, E. Adeli, M. G. Linguraru, Eds. (Springer International Publishing, Cham, 2019), Lecture Notes in Computer Science, pp. 167–175.

48. R. A. Kievit, S. W. Davis, J. Griffiths, M. M. Correia, Cam-CAN, R. N. Henson, A watershed model of individual differences in fluid intelligence. Neuropsychologia. 91, 186–198 (2016).

49. J. Dubois, P. Galdi, Y. Han, L. K. Paul, R. Adolphs, Resting-State Functional Brain Connectivity Best Predicts the Personality Dimension of Openness to Experience. Personal. Neurosci. 1 (2018), doi:10.1017/pen.2018.8.

50. W.-T. Hsu, M. D. Rosenberg, D. Scheinost, R. T. Constable, M. M. Chun, Resting-state functional connectivity predicts neuroticism and extraversion in novel individuals. Soc. Cogn. Affect. Neurosci. 13, 224–232 (2018).

51. A. D. Nostro, V. I. Müller, D. P. Varikuti, R. N. Pläschke, F. Hoffstaedter, R. Langner, K. R. Patil, S. B. Eickhoff, Predicting personality from network-based resting-state functional connectivity. Brain Struct. Funct. 223, 2699–2719 (2018).

52. U. Pervaiz, D. Vidaurre, M. W. Woolrich, S. M. Smith, Optimising network modelling methods for fMRI. NeuroImage. 211, 116604 (2020).

53. A. S. Greene, S. Gao, D. Scheinost, R. T. Constable, Task-induced brain state manipulation improves prediction of individual traits. Nat. Commun. 9, 1–13 (2018).

54. M. L. Elliott, A. R. Knodt, M. Cooke, M. J. Kim, T. R. Melzer, R. Keenan, D. Ireland, S. Ramrakha, R. Poulton, A. Caspi, T. E. Moffitt, A. R. Hariri, General functional connectivity: Shared features of resting-state and task fMRI drive reliable and heritable individual differences in functional brain networks. NeuroImage. 189, 516–532 (2019).

55. R. Jiang, N. Zuo, J. M. Ford, S. Qi, D. Zhi, C. Zhuo, Y. Xu, Z. Fu, J. Bustillo, J. A. Turner, V D. Calhoun, J. Sui, Task-induced brain connectivity promotes the detection of individual differences in brain-behavior relationships. NeuroImage. 207, 116370 (2020).

56. N. Raz, U. Lindenberger, P. Ghisletta, K. M. Rodrigue, K. M. Kennedy, J. D. Acker, Neuroanatomical Correlates of Fluid Intelligence in Healthy Adults and Persons with Vascular Risk Factors. Cereb. Cortex. 18, 718–726 (2008).

57. J. A. Amat, R. Bansal, R. Whiteman, R. Haggerty, J. Royal, B. S. Peterson, Correlates of intellectual ability with morphology of the hippocampus and amygdala in healthy adults. Brain Cogn. 66, 105–114 (2008).

58. M. S. Oechslin, D. Van De Ville, F. Lazeyras, C.-A. Hauert, C. E. James, Degree of Musical Expertise Modulates Higher Order Brain Functioning. Cereb. Cortex. 23, 2213–2224 (2013).

59. B. Zhu, C. Chen, X. Dang, Q. Dong, C. Lin, Hippocampal subfields’ volumes are more relevant to fluid intelligence than verbal working memory. Intelligence. 61, 169–175 (2017).

60. R. Li, J. Zhang, X. Wu, X. Wen, B. Han, Brain-wide resting-state connectivity regulation by the hippocampus and medial prefrontal cortex is associated with fluid intelligence. Brain Struct. Funct. 225, 1587–1600 (2020).

61. C. McNulty, thesis (2020).

62. H. C. Breiter, R. L. Gollub, R. M. Weisskoff, D. N. Kennedy, N. Makris, J. D. Berke, J. M. Goodman, H. L. Kantor, D. R. Gastfriend, J. P. Riorden, R. T. Mathew, B. R. Rosen, S. E. Hyman, Acute Effects of Cocaine on Human Brain Activity and Emotion. Neuron. 19, 591–611 (1997).

63. I. Aharon, N. Etcoff, D. Ariely, C. F. Chabris, E. O’Connor, H. C. Breiter, Beautiful Faces Have Variable Reward Value: fMRI and Behavioral Evidence. Neuron. 32, 537–551 (2001).

64. J. M. Gilman, S. Lee, J. K. Kuster, M. J. Lee, B. W. Kim, A. van der Kouwe, A. J. Blood, H. C. Breiter, Variable activation in striatal subregions across components of a social influence task in young adult cannabis users. Brain Behav. 6, e00459 (2016).

65. H. C. Breiter, I. Aharon, D. Kahneman, A. Dale, P Shizgal, Functional Imaging of Neural Responses to Expectancy and Experience of Monetary Gains and Losses. Neuron. 30, 619–639 (2001).

66. H. C. Breiter, G. P. Gasic, N. Makris, in Complex Systems Science in Biomedicine, T. S. Deisboeck, J. Y Kresh, Eds. (Springer US, Boston, MA, 2006; http://link.springer.com/10.1007/978-0-387-33532-2_33), Topics in Biomedical Engineering International Book Series, pp. 763–810.

67. E. J. Nestler, S. E. Hyman, D. M. Holtzman, R. C. Malenka, Molecular Neuropharmacology: A Foundation for Clinical Neuroscience (McGraw-Hill Education, New York, NY, ed. 3, 2015; neurology. mhmedical.com/content.aspx? aid=1105917072).

68. F. Nemmi, C. Nymberg, E. Helander, T. Klingberg, Grit Is Associated with Structure of Nucleus Accumbens and Gains in Cognitive Training. J. Cogn. Neurosci. 28, 1688–1699 (2016).

69. H. C. Breiter, N. L. Etcoff, P. J. Whalen, W. A. Kennedy, S. L. Rauch, R. L. Buckner, M. M. Strauss, S. E. Hyman, B. R. Rosen, Response and Habituation of the Human Amygdala during Visual Processing of Facial Expression. Neuron. 17, 875–887 (1996).

70. A. K. Barbey, R. Colom, E. J. Paul, J. Grafman, Architecture of fluid intelligence and working memory revealed by lesion mapping. Brain Struct. Funct. 2, 485–494 (2014).

71. P.-Y. Chen, C.-L. Chen, Y.-C. Hsu, W.-Y. I. Tseng, Fluid intelligence is associated with cortical volume and white matter tract integrity within multiple-demand system across adult lifespan. NeuroImage. 212, 116576 (2020).

72. Y. Y. Choi, N. A. Shamosh, S. H. Cho, C. G. DeYoung, M. J. Lee, J.-M. Lee, S. I. Kim, Z.-H. Cho, K. Kim, J. R. Gray, K. H. Lee, Multiple Bases of Human Intelligence Revealed by Cortical Thickness and Neural Activation. J. Neurosci. 28, 10323–10329 (2008).

73. F. J. Román, F. J. Abad, S. Escorial, M. Burgaleta, K. Martínez, J. Álvarez-Linera, M. Á. Quiroga, S. Karama, R. J. Haier, R. Colom, Reversed hierarchy in the brain for general and specific cognitive abilities: A morphometric analysis. Hum. Brain Mapp. 35, 3805–3818 (2014).

74. E. Vuoksimaa, M. S. Panizzon, C.-H. Chen, M. Fiecas, L. T. Eyler, C. Fennema-Notestine, D. J. Hagler, B. Fischl, C. E. Franz, A. Jak, M. J. Lyons, M. C. Neale, D. A. Rinker, W. K. Thompson, M. T. Tsuang, A. M. Dale, W. S. Kremen, The Genetic Association Between Neocortical Volume and General Cognitive Ability Is Driven by Global Surface Area Rather Than Thickness. Cereb. Cortex. 25, 2127–2137 (2015).

75. R. A. Kievit, D. Fuhrmann, G. S. Borgeest, I. L. Simpson-Kent, R. N. A. Henson, The neural determinants of age-related changes in fluid intelligence: a pre-registered, longitudinal analysis in UK Biobank. Wellcome Open Res. 3 (2018), doi:10.12688/wellcomeopenres.14241.2.

76. S. Frangou, X. Chitins, S. C. R. Williams, Mapping IQ and gray matter density in healthy young people. NeuroImage. 23, 800–805 (2004).

77. M. M. Pangelinan, G. Zhang, J. W. VanMeter, J. E. Clark, B. D. Hatfield, A. J. Haufler, Beyond age and gender: Relationships between cortical and subcortical brain volume and cognitive-motor abilities in school-age children. NeuroImage. 54, 3093–3100 (2011).

78. D. Li, T. Liu, X. Zhang, M. Wang, D. Wang, J. Shi, Fluid intelligence, emotional intelligence, and the Iowa Gambling Task in children. Intelligence. 62, 167–174 (2017).

79. S. A. Bunge, N. M. Dudukovic, M. E. Thomason, C. J. Vaidya, J. D. E. Gabrieli, Immature Frontal Lobe Contributions to Cognitive Control in Children: Evidence from fMRI. Neuron. 33, 301–311 (2002).

80. J. M. Court, Immature brain in adolescence. J. Paediatr. Child Health. 49, 883–886 (2013).

81. D. M. Barch, M. D. Albaugh, S. Avenevoli, L. Chang, D. B. Clark, M. D. Glantz, J. J. Hudziak, T. L. Jernigan, S. F. Tapert, D. Yurgelun-Todd, N. Alia-Klein, A. S. Potter, M. P. Paulus, D. Prouty, R. A. Zucker, K. J. Sher, Demographic, physical and mental health assessments in the adolescent brain and cognitive development study: Rationale and description. Dev. Cogn. Neurosci. 32, 55–66 (2018).

82. D. C. Van Essen, K. Ugurbil, E. Auerbach, D. Barch, T. E. J. Behrens, R. Bucholz, A. Chang, L. Chen, M. Corbetta, S. W. Curtiss, S. Della Penna, D. Feinberg, M. F. Glasser, N. Harel, A. C. Heath, L. Larson-Prior, D. Marcus, G. Michalareas, S. Moeller, R. Oostenveld, S. E. Petersen, F. Prior, B. L. Schlaggar, S. M. Smith, A. Z. Snyder, J. Xu, E. Yacoub, The Human Connectome Project: A data acquisition perspective. NeuroImage. 62, 2222–2231 (2012).

83. E. A. Aeschlimann, A. E. Voelke, C. M. Roebers, Short-Term Storage and Executive Working Memory Processing Predict Fluid Intelligence in Primary School Children. J. Intell. 5, 17 (2017).

84. C. A. Sandman, K. Head, L. T. Muftuler, L. Su, C. Buss, E. Poggi. Davis, Shape of the basal ganglia in preadolescent children is associated with cognitive performance. NeuroImage. 99, 93–102 (2014).

85. A. Abedelahi, H. Hasanzadeh, H. Hadizadeh, M. T. Joghataie, Morphometric and volumetric study of caudate and putamen nuclei in normal individuals by MRI: Effect of normal aging, gender and hemispheric differences. Pol. J. Radiol. 78, 7–14 (2013).

86. M.-M. Mesulam, Principles of Behavioral and Cognitive Neurology (Oxford University Press, 2000).

87. S. N. Haber, Corticostriatal circuitry. Dialogues Clin. Neurosci. 18, 7–21 (2016).

88. A. D. Lawrence, B. J. Sahakian, T. W. Robbins, Cognitive functions and corticostriatal circuits: insights from Huntington’s disease. Trends Cogn. Sci. 2, 379–388 (1998).

89. J. Brand, F. W. Bylsma, E. H. Aylward, J. Rothlind, C. A. Gow, Impaired source memory in huntington’s disease and its relation to basal ganglia atrophy. J. Clin. Exp. Neuropsychol. 17, 868–877 (1995).

90. L. C. Dang, G. R. Samanez-Larkin, J. S. Young, R. L. Cowan, R. M. Kessler, D. H. Zald, Caudate asymmetry is related to attentional impulsivity and an objective measure of ADHD-like attentional problems in healthy adults. Brain Struct. Funct. 221, 277–286 (2016).

91. B. J. Casey, T. Cannonier, M. I. Conley, A. O. Cohen, D. M. Barch, M. M. Heitzeg, M. E. Soules, T. Teslovich, D. V. Dellarco, H. Garavan, C. A. Orr, T. D. Wager, M. T. Banich, N. K. Speer, M. T. Sutherland, M. C. Riedel, A. S. Dick, J. M. Bjork, K. M. Thomas, B. Chaarani, M. H. Mejia, D. J. Hagler, M. Daniela Cornejo, C. S. Sicat, M. P. Harms, N. U. F. Dosenbach, M. Rosenberg, E. Earl, H. Bartsch, R. Watts, J. R. Polimeni, J. M. Kuperman, D. A. Fair, A. M. Dale, The Adolescent Brain Cognitive Development (ABCD) study: Imaging acquisition across 21 sites. Dev. Cogn. Neurosci. 32, 43–54 (2018).

92. D. C. Van Essen, S. M. Smith, D. M. Barch, T. E. J. Behrens, E. Yacoub, K. Ugurbil, The WU-Minn Human Connectome Project: An overview. NeuroImage. 80, 62–79 (2013).

93. N. Akshoomoff, J. L. Beaumont, P. J. Bauer, S. S. Dikmen, R. C. Gershon, D. Mungas, J. Slotkin, D. Tulsky, S. Weintraub, P. D. Zelazo, R. K. Heaton, Viii. Nih Toolbox Cognition Battery (cb): Composite Scores of Crystallized, Fluid, and Overall Cognition. Monogr. Soc. Res. Child Dev. 78, 119–132 (2013).

94. P. Besson, R. Lopes, X. Leclerc, P. Derambure, L. Tyvaert, Intra-subject reliability of the high-resolution whole-brain structural connectome. NeuroImage. 102 Pt 2, 283–293 (2014).

95. M. Defferrard, X. Bresson, P. Vandergheynst, in Advances in Neural Information Processing Systems 29, D. D. Lee, M. Sugiyama, U. V. Luxburg, I. Guyon, R. Garnett, Eds. (Curran Associates, Inc., 2016; http://papers.nips.cc/paper/6081-convolutional-neural-networks-on-graphs-with-fast-localized-spectral-filtering.pdf), pp. 3844–3852.

96. K. He, X. Zhang, S. Ren, J. Sun, in 2016 IEEE Conference on Computer Vision and Pattern Recognition (CVPR) (IEEE, Las Vegas, NV, USA, 2016; http://ieeexplore.ieee.org/document/7780459/), pp. 770–778.

97. A. Veit, M. J. Wilber, S. Belongie, in Advances in Neural Information Processing Systems 29, D. D. Lee, M. Sugiyama, U. V. Luxburg, I. Guyon, R. Garnett, Eds. (Curran Associates, Inc., 2016; http://papers.nips.cc/paper/6556-residual-networks-behave-like-ensembles-of-relatively-shallow-networks.pdf), pp. 550–558.

98. H. Liang, F. Zhang, X. Niu, Investigating systematic bias in brain age estimation with application to post-traumatic stress disorders. Hum. Brain Mapp. 40, 3143–3152 (2019).

99. R. R. Selvaraju, M. Cogswell, A. Das, R. Vedantam, D. Parikh, D. Batra, Grad-CAM: Visual Explanations from Deep Networks via Gradient-based Localization. Int. J. Comput. Vis. (2019), doi:10.1007/s11263-019-01228-7.

100. U. von Luxburg, A tutorial on spectral clustering. Stat. Comput. 17, 395–416 (2007).

101. M. Rubinov, O. Sporns, Complex network measures of brain connectivity: Uses and interpretations. NeuroImage. 52, 1059–1069 (2010).

